# Feeding-state dependent neuropeptidergic modulation of reciprocally interconnected inhibitory neurons biases sensorimotor decisions in *Drosophila*

**DOI:** 10.1101/2023.12.26.573306

**Authors:** Eloïse de Tredern, Dylan Manceau, Alexandre Blanc, Panagiotis Sakagiannis, Chloe Barre, Victoria Sus, Francesca Viscido, Md Amit Hasan, Sandra Autran, Martin Nawrot, Jean-Baptiste Masson, Tihana Jovanic

## Abstract

Animals’ feeding state changes behavioral priorities and thus influences even non-feeding related decisions. How is the feeding state information transmitted to non-feeding related circuits and what are the circuit mechanisms involved in biasing non-feeding related decisions remains an open question. By combining calcium imaging, neuronal manipulations, behavioral analysis and computational modeling, we determined that the competition between different aversive responses to mechanical cues is biased by feeding state changes. We found that this is achieved by differential modulation of two different types of reciprocally connected inhibitory neurons promoting opposing actions. This modulation results in a more frequent active type of response and less frequently a protective type of response if larvae are fed sugar compared to when they are fed a balanced diet. The information about the internal state is conveyed to the inhibitory neurons through homologues of the vertebrate neuropeptide Y known to be involved in regulating feeding behavior.

## Introduction

Physiological states like hunger and thirst are powerful regulators of behavior across the animal kingdom due to strong homeostatic drives that are critical for survival^1–7^. For example, across model systems, food-deprivation was shown to modulate responsiveness to stimuli by influencing sensory neurons and sensory pathways^8–11^, suggesting that food-deprivation can alter the perceived value of a stimulus, which in turn affects the behavioral decisions. Various studies have implicated changes in central processing^12–15^ which lead to changes in behavioral decisions in hungry animals. However, the detailed neural circuit mechanisms of this state-dependent flexibility of behaviors remain largely unknown.

Internal drives (e.g., hunger and thirst) need to be balanced by environmental demands such as the need to avoid dangers. Avoiding dangers is a critical instinctive behavior that needs to be balanced with finding and consuming food in order to ensure survival. Avoidance behaviors tend to be robust, which makes them excellent systems for studying neural bases of behavior^16–19^; yet they also need to be flexible in order for animals to adapt their behavioral strategies to different contexts and according to different internal states^17,20–22^. Feeding states and contexts can for example influence both the tolerance to the level of threat and action selection during threat avoidance^16–18,23^.

At the neural circuit level, such behavioral flexibility is thought to be implemented by neuromodulation (modulation of existing synaptic connections by neuropeptides, for instance) that could bias the outcome of competition between diverse behaviors. The outcome could in that case differ depending on the neuropeptide released^24–30^. Alternatively, information rerouting (using alternative circuit pathways)^12,13,31^ where information is processed differently depending on the context or state, could alter behavior choice in a context-dependent manner. The types of circuit motifs underlying competitive selection must allow for such flexible processing of information. However, the detailed neural circuit mechanisms underlying the state-dependent modulation of behavior, the neuromodulators involved, and their mechanism of actions on specific circuits, especially those that pertain directly to non-feeding or non-water seeking behavior, are not well understood.

This paper addresses the understudied aspect of the influence of physiological drives on competitive selection during avoidance behaviors and investigates the neural circuit mechanisms involved in a powerful model organism for neural circuit analysis: the *Drosophila* larva^32^. *Drosophila* larvae are ideally suited for combining comprehensive, synaptic-resolution circuit mapping in electron microscopy (EM) across the nervous system^33–36^ with targeted manipulation of uniquely identified circuit motifs at the individual neuron level, which makes it possible to establish causal relationships between circuit structure and function in a brain-wide manner. In addition, evolutionarily conserved neuropeptidergic and hormonal pathways in *Drosophila* have been shown to regulate its diverse behaviors^11,18,27,37,38^.

Previous studies have described larval avoidance response to a mechanical stimulus (air-puff) and detailed the neural circuit underlying the competition between “Hunching”, a protective or “startle-like” type of behavior, and “Head Casting”, an “active” exploratory type of behavior that can lead to escape^35,39–43^. We identified circuit motifs underlying competitive interactions between behavioral actions (reciprocal inhibition of inhibition) and sequence transitions (lateral and feedback disinhibition) between the two behaviors. These types of motifs based on disinhibition would allow for flexible behavioral selection in a context/state dependent manner. By combining calcium imaging, neuronal and neuropeptide manipulation at the single cell level with automated tracking, behavior classification and computational modeling, this work shows that larval responses to the air-puff are biased towards less protective actions and towards more active, exploratory actions upon changes in their feeding condition. We determine that this bias is due to the differential modulation of two reciprocally connected inhibitory neurons that drive competing behaviors (Hunching and Head Casting): the activity of the neuron that promotes the protective Hunching is decreased and the activity of the neuron that inhibits Hunching is increased. We also show that the modulation at the level of reciprocally interconnected inhibitory neurons results in a bias at the level of the network output, towards a state that will lead to Head Casting at the expense of Hunching upon feeding on sucrose. Finally, we determine that NPF and sNPF modulate the activity of the reciprocally interconnected inhibitory neurons to bias the behavioral output in a feeding state-dependent manner.

## RESULTS

### Changes in feeding conditions for short periods of time affect larval feeding and locomotion

To study the effect of diet on behavioral decisions as a result of cognitive control and prioritization of needs and motivations rather than to long-term physiological changes that would affect circuit properties, we sought to establish a food-deprivation protocol of short duration. We determined the shortest duration of food deprivation that was sufficient to induce quantifiable changes in behavior. Depriving larvae of food completely (by putting them on a water-soaked filter paper) or feeding them on 20% sucrose only (and therefore depriving them of proteins and increasing their sugar intake) for 90 mins was sufficient to alter larval locomotion. Starved larvae and larvae fed on sucrose moved at a higher speed and spent more time Crawling than normally-fed larvae, eventually dispersing faster in less-curved trajectories (Fig. 1D, Extended Data Fig. 1A-B). These changes in locomotion are consistent with increased exploration and are likely due to an increased drive to find nutrients caused by deprivation. In order to determine that the 90 minutes feeding deprivation protocol induced changes in behavior that were reversible, we monitored locomotion in these larvae upon refeeding them for 15 minutes. Upon refeeding, larvae were slower and their locomotion phenotype was similar to the one of larvae that were constantly fed (Extended Data Fig. 1C).

**Fig. 1.**
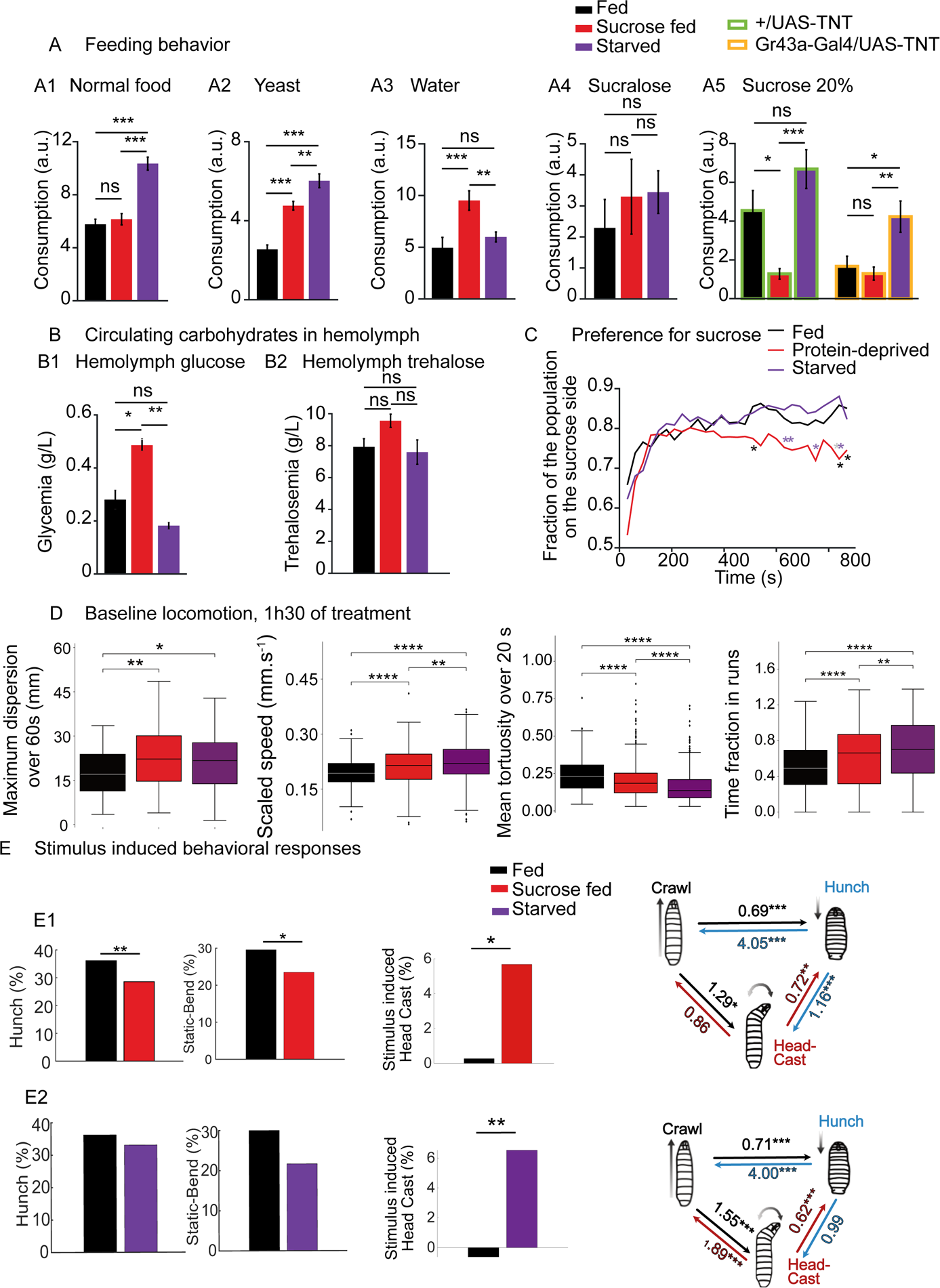
Changes in physiology and behavior of food-deprived larvae. A: Larval feeding on different substrates was quantified, in control animals fed ad-libitum, in animals fed on sucrose or in animals subjected to complete starvation during 90 min. A1: Starved animals increased their feeding on a standard food medium as compared to fed animals, while animals fed on sucrose only did not (ANOVA, n = 35-41 larvae, Tukey post-hoc test ***: p < 0.001). A2: Both animals fed on sucrose only and starved animals increase their yeast feeding as compared to normally fed larvae (ANOVA, n = 51-59 larvae, Tukey post-hoc test ***: p < 0.001, **: p < 0.01). A3: While starved animals consume a similar amount of water as fed ones, animals fed on sucrose only double their water consumption as compared to fed and starved larvae (ANOVA, n = 52-54 larvae, Tukey post-hoc test ***: p < 0.001, **: p < 0.01). A4: Consumption of the non-energetic sweetener sucralose was similar among conditions (n = 25-32 larvae). A5: Larvae that had fed on a 20% sucrose solution for 90 min decreased their sucrose intake as compared to fed and starved larvae (ANOVA, n = 29-33 larvae, Tukey post-hoc test ***: p < 0.001, *: p < 0.05). Gustatory neurons inhibition (Gr43a-Gal4>TNT) decreases sucrose intake in fed larvae to the level of larvae fed on sucrose, while the consumption of starved larvae remains higher (ANOVA, n = 24-33 larvae, Tukey post-hoc test **: p < 0.01, *: p < 0.05). B: Concentration of glucose and trehalose in larval hemolymph. B1: Glucose concentration is increased in the hemolymph of larvae that were fed on 20% sucrose solution for 90 min compared to larvae fed on a standard diet and to starved animals (ANOVA, N = 3 samples from 5 larvae each, Tukey post-hoc test **: p < 0.01, *: p < 0.05). B2: Trehalose concentration is similar in the hemolymph of larvae fed on the different diets (N = 6 samples from 10 larvae each). C: Place preference assay for sucrose. Larvae that fed on a 20% sucrose solution for 90 min exhibit a decreased glucose preference compared to normally fed and starved larvae (n = 79-133 larvae, Chi-2 test **: p < 0.01, *: p < 0.05). D: Analysis of larval locomotion in the absence of sensory stimulus. After 90 min of sucrose feeding or complete starvation, larval dispersion, time spent crawling and speed are significantly increased, while the tortuosity of the trajectory is decreased compared to larvae fed on a standard diet (Mann-Whitney test, ****: p < 0.0001, ***: p < 0.001, **: p < 0.01, *: p < 0.05). E: Manipulating the feeding state modulates behavioral responses to mechanosensory stimuli. Behavior in response to air-puff during the first five seconds uoin stimulus onset, for, E1 in larvae fed on sucrose only (n = 592-629 larvae) and E2 starved larvae compared to larvae fed on standard food (n =704-771 larvae, ***: p < 0.001, **: p < 0.01, *: p < 0.05). See also Extended Data Fig. 1 and Supplementary Tables 2, 3, 4 and 5.

To determine whether the larval need for nutrients was affected by the feeding protocols that they were subjected to, the consumption rate of different foods was quantified in different feeding conditions. Starved larvae significantly increased their intake of standard *Drosophila* food and yeast (rich in amino acids) compared to both normally-fed larvae and larvae that were fed on sucrose (Fig. 1A), which is consistent with a deficit in nutrients in starved larvae. Larvae fed on sucrose significantly increased their intake of yeast, which is consistent with a deficit in amino acids (Fig. 1A). These larvae, however, did not increase their intake of standard *Drosophila* food, suggesting that the increased sugar consumption might suppress their intake of carbohydrate-rich food. To test whether larvae fed on sucrose were repulsed by sugar, a choice assay was performed: larvae were added to the middle of an agar plate where half of the plate was covered in agar and the other half in agar mixed with sucrose. The larvae were then monitored for 15 minutes (Fig. 1C). After the 15 minutes, more larvae were found on the agar+sucrose half of the arena in normally-fed and starved larvae, with a preference index increasing over time, while larvae fed on sucrose showed a decrease in preference for agar supplemented with sucrose. Larvae fed on only 20% sucrose for 90 minutes significantly decreased their sucrose intake compared to fed and starved larvae, and their sucralose intake was similar to that of normally-fed and starved larvae (Fig. 1A4), suggesting that their avoidance of sugar was mediated by energy sensing pathways and not taste.

To determine whether the 90-minute food-deprivation or sucrose diet protocols caused changes in glucose levels, the hemolymph glucose levels were quantified in the different feeding conditions. Glucose levels were slightly but not significantly decreased in starved larvae compared to the fed larvae. In larvae that were fed on 20% sucrose, the glucose level was significantly increased compared to both normally-fed and starved larvae (Fig. 1B). This high glucose level suggests that larvae fed on sucrose suppress consumption of carbohydrates due to high circulating levels of glucose. Finally, the water consumption of larvae in different feeding conditions was quantified. Larvae fed on 20% sucrose increased their water consumption significantly compared to both normally-fed and starved larvae (Fig 1A3), likely due to changes in extracellular osmolality as a result of high sugar intake^6^. We therefore tested whether rehydration would reduce the increase in exploration and locomotion observed in larvae fed on sucrose. Indeed, after larvae fed on sucrose only were put on water for 15 minutes, their locomotion was similar to the ones of normally fed larvae (Extended Data Fig. 1D).

Altogether, these results suggest that depriving larvae of nutrients for 90 minutes is sufficient to alter feeding and larval locomotion and that these changes are due to lack of nutrients and lower energy levels in starved larvae and protein hunger and thirst in larvae fed on sucrose. The observed increase in locomotion in larvae deprived of nutrients would thus likely result from an increase in motivational drive to find the missing nutrients.

### Changes in physiological states affect sensorimotor decisions in response to an air-puff

To determine whether starvation and a sucrose-only diet can affect non-feeding-related behaviors, we monitored *Drosophila* larva sensorimotor decisions in response to an aversive mechanical stimulus, the air-puff. In response to an air-puff, larvae perform probabilistic sequences of five mutually exclusive actions that we have characterized in detail in the past^35,44^: Hunch, Bend, Stop, Back-up, and Crawl. We have also identified circuit motifs and characterized the neural circuit mechanisms underlying the competitive interactions between the two most prominent actions (i.e. the Hunch and the Bend) that occur in response to an air-puff^35^. The model in that study predicts that the different activation levels of inhibitory neurons determine which actions will take place: the Hunch, Bend, or the Hunch Bend sequence.

With characterized neurons and synaptic connectivity between the neurons, as well as the availability of driver lines that label neurons of interest, this circuit provides an excellent system to study whether the activity of the neurons is modulated by the changes in an animal’s physiological state.

To determine whether the feeding state modulates larval behavior in response to the mechanosensory stimulus and, by extension, whether sensorimotor decision-making in response to an air-puff is an adequate behavioral paradigm to study the effects of food deprivation on neural circuit activity, we compared larval behavioral responses to an air-puff after subjecting them to the different feeding protocols (as described in the first section of the Results). For this purpose, we used automated tracking^45^ to monitor larval behavior in response to an air-puff and updated the machine-learning-based classification method developed in our previous work^44^ to compute probabilities of the different actions that occur in response to an air-puff. The new classifiers were trained on larvae fed on different diets, thus taking into account a broader range of behavioral dynamics, and separated the bend behavioral category into two types of bending behavior: Static Bends, a form of protective action where the larva responds to the stimulus by immobilizing in a curved position for a period of time, and the exploratory active Head Casts that can lead to escape^39^.

Our analysis showed that larvae fed on sucrose Hunch less (and perform less of Static Bends) and Head Cast more (Fig. 1E1). Similarly, starved larvae performed less Static Bends and more Head Casts (Fig. 1E2). Transitions to Hunching were decreased in larvae fed on sucrose and starved larvae, while transitions to Head Cast were increased (from Hunching to Head Casting in larvae fed on sucrose only and from Crawl to Head Casting in both starved larvae and larvae fed on sucrose) (Fig. 1E). Increasing duration of sucrose feeding and starvation to 5h resulted in similar phenotypes (Extended Data Fig. 1E, F).

Using our machine-learning based classification to monitor internal state induced behavioral changes revealed that changing feeding conditions for short periods of time alters sensorimotor decisions in response to an aversive mechanical cue and results in less protective type of actions, and more actions consistent with active exploration and escape.

### The feeding state does not affect the responses of chordotonal sensory neurons to a mechanical stimulus

To test whether changes in feeding conditions alter the perceived value of a stimulus, which in turn would affect the behavioral decisions, calcium responses in sensory neurons that sense the air-puff, the chordotonal sensory neurons^35,41,44,45^ were monitored in response to a moderately strong mechanical stimulus in the three different feeding conditions: normally fed, fed on sucrose and starved (Extended Data Fig. 2B). We found no significant differences in chordotonal calcium responses in the different feeding states (Fig. 2B, Extended Data Fig. 2A), although the responses were slightly higher in starved larvae compared to the larvae fed on standard food and on sucrose only. To ensure that the changes in feeding conditions did not influence protein expression and, by extension, GCAMP expression levels, GFP was genetically expressed in the neurons and its expression level was quantified by comparing the fluorescence in larvae exposed to all three feeding conditions. No differences were found in GFP fluorescence in the three feeding states (Extended Data Fig. 2C). The results therefore suggest that chordotonal responses are not significantly affected by starvation or sucrose feeding and that the behavioral changes observed in the different feeding conditions are not due to the changes in stimulus sensitivity at the level of sensory neurons. We further confirmed this by optogenetically activating chordotonal sensory neurons in larvae in all three feeding states using a driver that labels all eight subtypes of chordotonal sensory neurons: R61D08. We found that although the activation of chordotonal sensory neurons in all three feeding states was the same due to optogenetic stimulation, larvae performed fewer Hunches and more Head Casts when they were fed only on sucrose than when they were normally fed (Fig. 2C). Similarly, the modulation of behavior when they were starved could be observed upon chordotonal optogenetic activation (Extended Data Fig. 2D). Altogether, these results show that the feeding state-dependent modulation could target neurons downstream of the chordotonal sensory neurons.

**Fig. 2.**
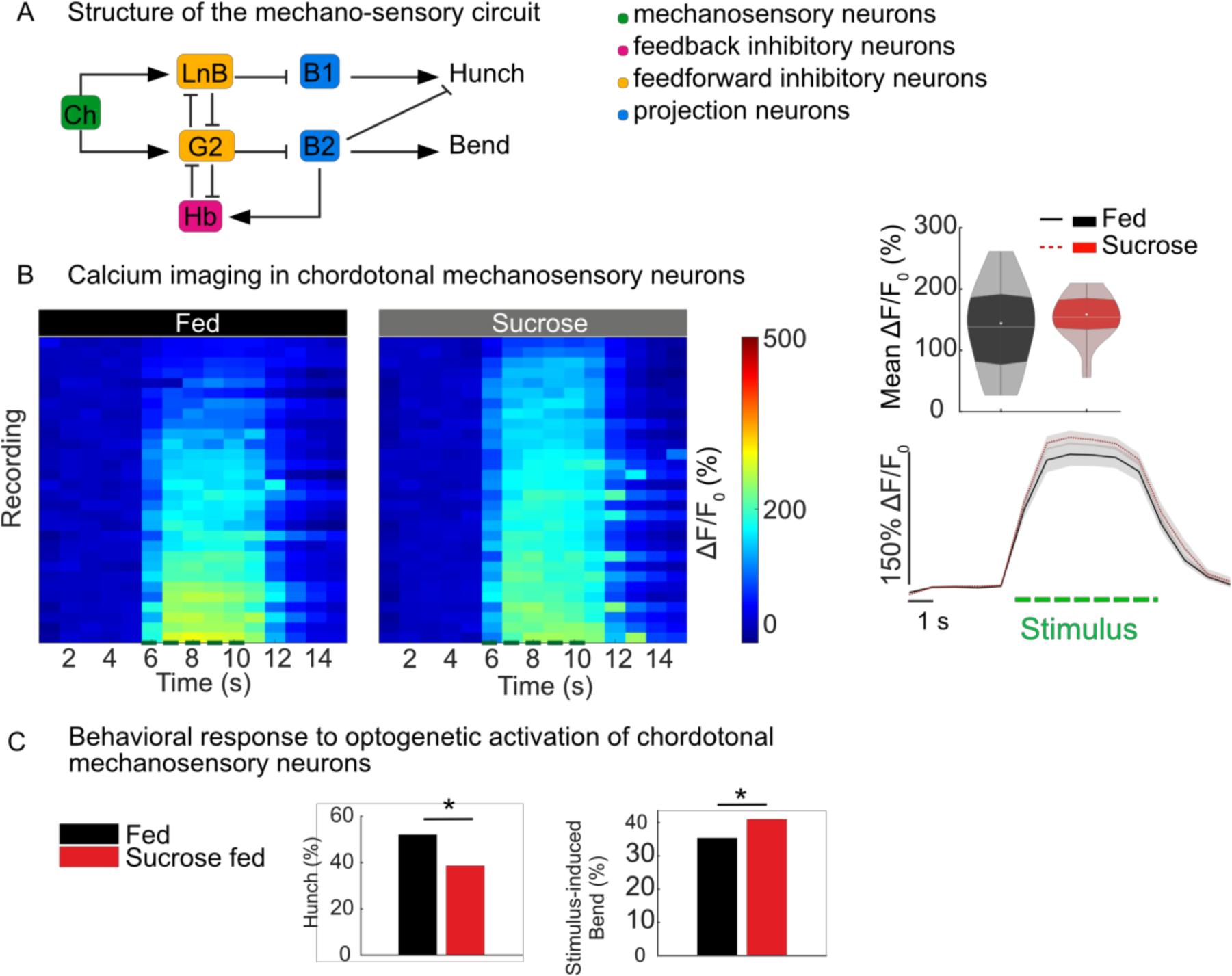
Feeding state dependent bias in sensorimotor responses does not come from the modulation of sensory neurons. A: Organization of the circuit. B: Calcium responses to mechanical stimulations in chordotonal neurons larvae fed on sucrose only and on standard food (R61D08-Gal4/UAS-GCaMP6s). Left panel: calcium responses of chordotonal neuron projections in the VNC from different individuals fed on different diets. Lower right panel: mean calcium response of chordotonal neurons over time. The green dashed line corresponds to stimulus duration. Upper right panel: calcium response averaged during the stimulus. White line represents the mean, white dot represents the median. Stimulus-induced activity of chordotonal mechanosensory neurons is unchanged in animals fed on sucrose as compared to larvae fed on standard food (n = 10 larvae, 3 trials per larva, t-test p = 0.249). C: Optogenetic activation of cho in larvae fed only on sucrose. Hunch probability is computed during the first 2 seconds from stimulus onset. Bend is the mean probability during the first 10 seconds from stimulus onset, corrected by the baseline recording prior to the stimulus. (n = 192 larvae for fed, 168 for sucrose fed, p = 0.011 for Hunch, 0.028 for Bend). See also Extended Data Fig. 2 and Supplementary Tables 4 and 6.

### Feeding states modulate the output of the circuit for the choice between Hunch and Bend

To determine whether the changes in feeding state influence the output of the network for the selection between the Hunch and the Bend, characterized in previous work^35^, calcium responses of projection neurons Basin-1 and Basin-2 were monitored at different intensities of mechanical stimulation in the three feeding states. Basin-1 and Basin-2 are differentially involved in Hunching and Bending, as shown in Jovanic *et al*., 2016: Basin-1 is required for both Hunching and Bending, while Basin-2 promotes Bending and inhibits Hunching. Responses in Basin-2 neurons can thus be used as a read-out of the Bend state: if Basin-2 is ON, the larva will Bend, and if it is OFF, the larva will Hunch. We found that for most stimulus intensities, the Basin-1 responses remain only mildly affected by the sucrose-only diet (Fig. 3A, Extended Data Fig. 3C). Only for the very weak stimulus intensity do we observe a decrease in Basin-1 response (Extended Data Fig. 3C). In the starved condition, the effect on Basin-1 neurons was similar to the sucrose-only feeding condition (Extended Data Fig. 3A, C). The activity of Basin-2, on the other hand, was moderately increased in larvae fed only sucrose at all intensities except at the lowest stimulus intensity (Fig. 3B, Extended Data Fig. 3D), where there was no difference between Basin-2 responses in different feeding conditions (Extended Data Fig. 3D). However, the activity of Basin-2 in starved larvae was slightly (but not significantly) decreased compared to fed larvae at lower intensities of stimulation, whereas it was increased at higher intensities of stimulation (Extended Data Fig. 3B,D).

**Fig. 3.**
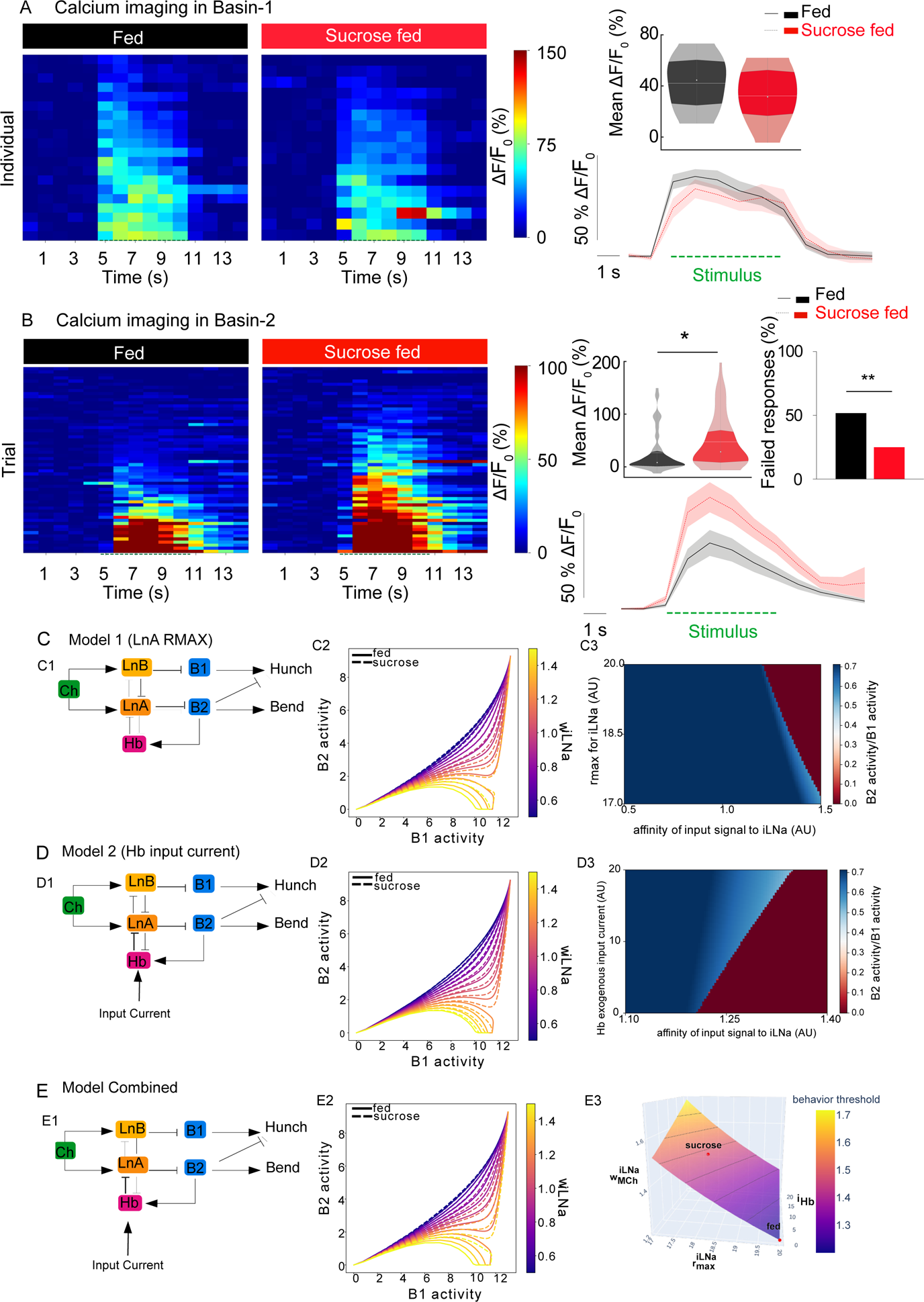
The feeding status affects the output of the sensorimotor circuit. A: Basin-1 calcium responses to mechanical stimulations in larvae fed on standard food and on sucrose only larvae (20B01-lexA; LexAop-GCaMP6s, UAS-CsChrimson-mCherry). Left panel: calcium responses of Basin-1 from different individuals fed on different diets. Lower right panel: mean calcium response of Basin-1 over time. The green dashed line corresponds to stimulus onset. Upper right panel: mean calcium response averaged during the stimulus. White line represents the mean, white dot represents the median. Stimulus-induced activity of Basin-1 neurons is similar in larvae fed on sucrose only compared to larvae fed on a standard food (n = 17/20 larvae, 1 trial per larva, t-test p = 0.2577). B: Basin-2 calcium responses to mechanical stimulations in different states (SS00739/UAS-GCaMP6s). Left panel: calcium responses of Basin-2 from different individuals fed on different diets. Lower right panel: mean calcium response of Basin-2 over time. The green dashed line corresponds to stimulus onset. Upper right panel: mean calcium response averaged during the stimulus. White line represents the mean, white dot represents the median. Low panel: percentage of trials that failed to elicit a calcium response in Basin-2. Stimulus-induced activity of Basin-2 neurons is enhanced in animals fed on sucrose only compared to larvae fed on standard food (n = 12 larvae, 5 trials per larva, mean response comparisons t-test p = 0.0118, percentage of failed responses comparison Chi-2 test p = 0.0027). C-E: Simple connectome-based rate model of the decision circuit for the selection Hunch and Bend/Hedcast C: Model 1: version of the model where the sucrose state is modeled by a decreased maximum rate of the LNa node. C1. schematic of the circuit where the influence of LNa on downstream targets is weaker C2. state space trajectories of Basin-1 (B1) and Basin-2 Activity as a function of coupling from MCh to LNa for model 1. C3. phase ratio phase diagram for model 1 D: Model 2: version of the model where the sucrose state is modeled by an input current to Hb) Handle B neuron D1. schematic of the circuit where the influence of Hb on downstream targets is stronger D2. state space trajectories of Basin-1 (B1) and Basin-2 Activity as a function of coupling from MCh to LNa for model 2. D3. phase ratio phase diagram for model 2 E: Combined model: version of the model where the sucrose state is modeled by both an input current to Handle B (Hb) neuron and decreased Maximum rate of LNa E1. schematic of the circuit where the influence of Hb on downstream targets is stronger and the one of Lna weaker E2. state space trajectories of Basin-1 (B1) and Basin-2 Activity as a function of coupling from MCh to LNa for the combined model. E3. behavior phase diagram for the combined model. See also Extended Data Fig. 3 and Extended Data Fig. 5 and Supplementary Table 6.

We then compared the distributions of Basin-2 responses in larvae fed on either standard food or sucrose to a moderately strong mechanical stimulus (5V applied to piezo at 1000 HZ)) and observed a higher probability of absence of neuronal response in normally-fed larvae (Extended Data Fig. 3E). The previous study has recorded depolarization responses simultaneously from Basin-1 and Basin-2 to a mechanical stimulus and has shown that Basin-1 neurons always respond while Basin-2 responses are probabilistic^35^. In this study, we computed the probability of Basin-2 responses to the mechanical stimulus and observed a significantly higher probability of responses that cross the threshold (see methods) in larvae fed on sucrose, which is consistent with higher Head Casting and lower Hunching probabilities in these larvae (Extended Data Fig. 3E). In starved larvae, similar trends were observed at higher stimulus intensities, while at medium intensity (5V), there was no difference between Basin-2 response probability in fed and starved larvae (Extended Data Fig. 3E). This difference in Basin-2 responses between larvae fed only sucrose and starved larvae could explain why Hunching is not consistently significantly reduced in starved larvae.

### Reciprocally connected Inhibitory interneurons in the decision circuit are differentially modulated by the feeding state

Based on the simulation of the model of the circuit for a choice between two air-puff-induced actions (Hunch and Bend)^35^, it is predicted that the relative level of activation of the reciprocally connected feedforward inhibitory interneurons determines the outcome of the competition: Basin-1 only state (Hunch) or Co-active state (Bend). In this study, we investigated if changes in the feeding state could affect the level of activation of two different classes of the inhibitory neurons: feedforward and feedback inhibitory neurons.

To address this, we started by building a simple rate model, that, as in the previous study^35^, is based on the observed connectivity between neuron classes from the EM reconstruction data, and where weights of excitatory or inhibitory connections between neurons were proportional to the number of synapses (Fig. 3C-E). As previously described, the model predicts that the level of activation of the two classes of feedforward inhibitory neurons will determine the output state of the network at the level of the Basin-1 and Basin-2 neurons (Fig. 3C-E, Extended Data Fig. 5A). The “sucrose state” was modeled in one version of the model as the decreased maximum intensity of LNa neuron activity to explore the role of the feedforward inhibitory neurons in the feeding state modulation (Fig. 3C). In order to explore the contribution of the feedback inhibitory neurons, in another version of the model, the sucrose state was represented by adding an input to the Handle-b neurons (Fig. 3D). Finally, we made a combined model where both the LNa and Handle-b were modulated in the sucrose state (Fig. 3E, Extended Data Fig. 5A-D). In all of the versions of the model the Hunching decreased and Head Casting increased in the sucrose state compared to normal fed state (Fig. 3C3, D3, E3).

These model results suggest that modulating both feedforward and feedback inhibitory neurons of the circuit could result in the observed behavioral changes upon sucrose feeding. To test this experimentally, calcium transients in response to a mechanical stimulus were imaged in the two different types of inhibitory neurons to which we had genetic access. An LNa neuron type, Griddle-2 (G2), that promotes Hunching^35^, showed significantly lower responses to a mechanical stimulus in larvae fed on sucrose compared to the normally fed larvae (Fig. 4A). The decrease in response was observed at different intensities of mechanical stimulation (Extended Data Fig. 4C). Similarly, in starved larvae, the responses of Griddle-2 neurons were also decreased across different intensities of stimulation (Extended Data Fig. 4A, C).

**Fig. 4.**
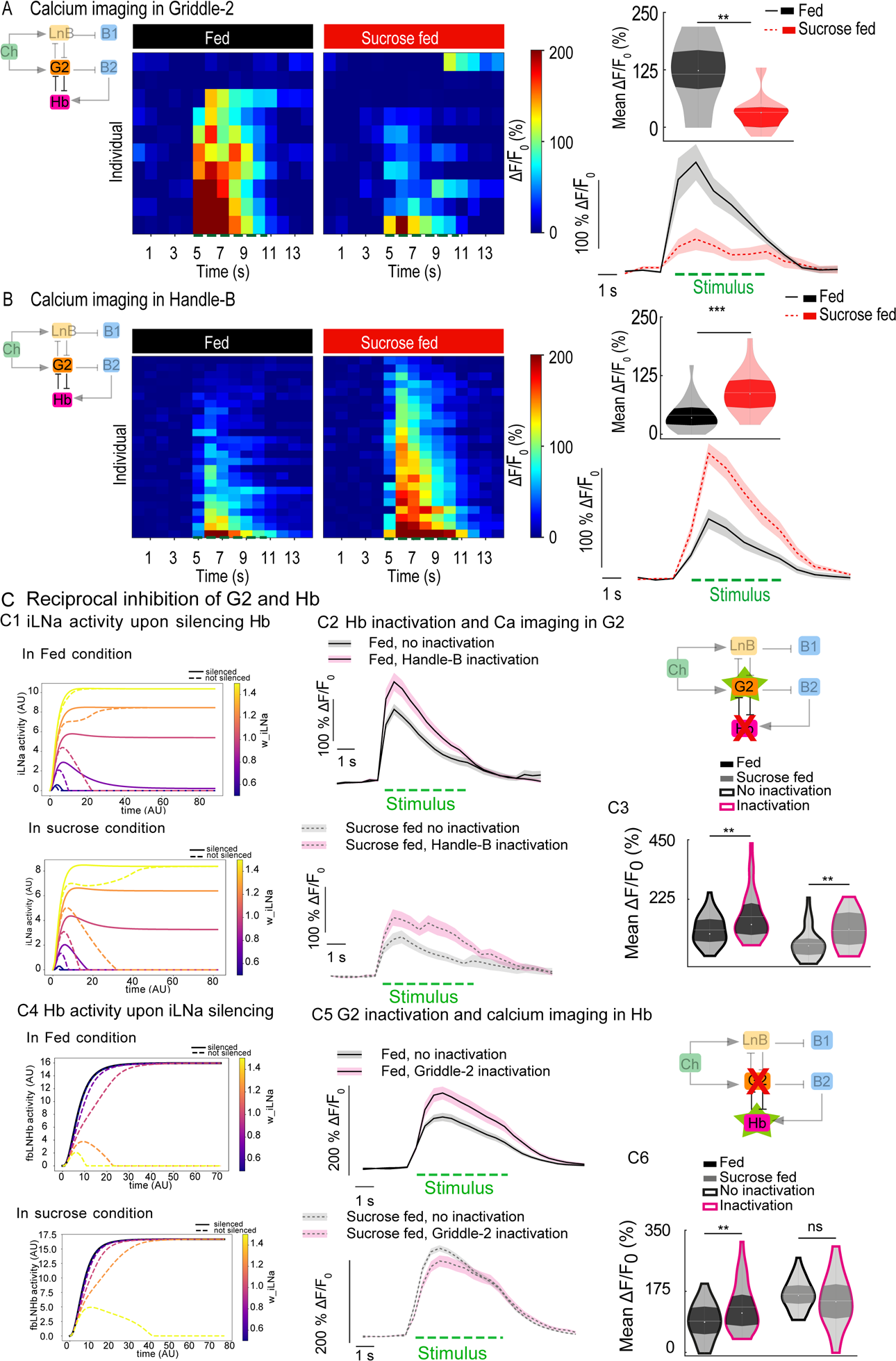
Two reciprocally connected interneurons are oppositely modulated by the feeding state. A: Griddle-2 calcium responses to mechanical stimulations in larvae fed on standard food and sucrose only (SS00918/UAS-GCaMP6s). Left panel: calcium responses of Griddle-2 from different individuals, fed on standard food or on sucrose only. Lower right panel: mean calcium response of Griddle-2 over time. The green dashed line corresponds to stimulus onset. Upper right panel: mean calcium response averaged during the stimulus. White line represents the mean, white dot represents the median. Stimulus-induced activity of Griddle-2 is decreased in animals fed on sucrose only compared to larvae fed on a standard food (n = 10 larvae, 1 trial per larva, t-test p = 0.0068). B: Handle-b calcium responses to mechanical stimulations in larvae fed on standard food or on sucrose only (SS00888/UAS-GCaMP6s). Left panel: calcium responses of Handle-b from different individuals, fed on standard food or on sucrose only. Lower right panel: mean calcium response of Handle-b over time. The green dashed line corresponds to stimulus onset. Upper right panel: mean calcium response averaged during the stimulus. White line represents the mean, white dot represents the median. Stimulus-induced activity of Handle-b is increased in animals fed on sucrose only compared to larvae fed on a standard food (n = 23/25 larvae, 1 trial per larva, t-test p = < 0.0001). C: Reciprocal inhibition between Handle-b and Griddle-2. C1: model simulation, trajectories of LNa activity upon Hb inactivation in fed (top panel) and sucrose condition (bottom panel) C2: Left panel: mean calcium response of Griddle-2 over time, with (SS0888-Gal4>UAS-TNT 55C05-LexA>LexAop-GCaMP6s) or without Handle-b inactivation (+/UAS-TNT 55C05-LexA>LexAop-GCaMP6s), in larvae fed on standard food or on sucrose only. The green dashed line corresponds to stimulus onset. C3: mean calcium response averaged during the stimulus. White line represents the mean, white dot represents the median. Griddle-2 responses to mechanical stimuli are increased after Handle-b inactivation, both in larvae fed on standard food (top panel) or on sucrose (bottom panel) (n = 7-9 larvae, 4 trials per larva, t-test after inactivation in larvae fed on standard food p = 0.009, fed on sucrose only p = 0.002). C1. model simulation, trajectories of Hb activity upon iLNa inactivation in fed (top panel) and sucrose condition (bottom panel) C5. mean calcium response of Handle-b over time, with (55C05-LexA>LexAop-TNT 22E09-Gal4>UAS-GCaMP6s) or without Handle-b inactivation (+/LexAop-TNT 22E09-Gal4>UAS-GCaMP6s), in larvae larvae fed on standard food or on sucrose. The green dashed line corresponds to stimulus onset. C6: mean calcium response averaged during the stimulus. White line represents the mean, white dot represents the median. Handle-b responses to mechanical stimuli are increased after Griddle-2 inactivation only in fed animals (n = 6-10 larvae, 5 trials per larva, t-test after inactivation in fed p = 0.0027, sucrose condition p = 0.065). See also Extended Data Fig. 4 and Extended Data Fig. 5 and Supplementary Table 6.

Based on the model, the decrease in the activity of LNa would result in less Hunching and more Head Casting which is consistent with the behavioral changes we observed after feeding on sucrose. The model has also shown that another motif in the circuit, feedback disinhibition, promotes Bending by amplifying the activity of Basin-2. We recorded Handle-b responses to the mechanical stimulus in the different feeding states and found that the Handle-b neurons, that inhibit Hunching^35^, show stronger responses to a mechanical stimulus in larvae fed on sucrose and in starved larvae compared to normally fed larvae (Fig. 4B, Extended Data Fig. 4B,D). This was true for all the different intensities of stimulation we tested (Extended Data Fig. 4D). Therefore, the decreased response of Griddle-2 neurons (Fig. 4A) and increased responses of Handle-b neurons (Fig. 4B) in larvae fed on sucrose could bias the behavioral outcome, consistent with the behavioral changes observed in larvae upon sucrose feeding (less Hunching and more Head Casting) (Fig. 1E1).

EM analysis shows that Griddle-2 and Handle-b neurons are reciprocally interconnected (Fig. 4A)^35^.The circuit and the model predict that silencing Handle-b results in an increase in Griddle-2 activity and vice-versa (Fig. 4C). Further, the reciprocal connections between Griddle-2 and Handle-b were probed functionally and the model predictions tested experimentally; calcium responses to a mechanical stimulus were monitored in Griddle-2 neurons while inactivating Handle-b using tetanus-toxin in both larvae fed on standard food and on sucrose only. The responses were increased compared to control larvae in both feeding states (Fig. 4C1-C3, Extended Data Fig. 4E), which is consistent with the Handle-b inhibition of Griddle-2. However, the increase in Griddle-2 responses upon Handle-b inactivation in larvae fed on sucrose did not reach the levels of Griddle-2 responses upon inactivation in larvae fed on standard food (Fig. 4C2-C3, Extended Data Fig. 4E). This could suggest that the increase in Handle-b activity after sucrose feeding is insufficient to reduce the Griddle-2 responses in larvae fed on sucrose and that both Handle-b and Griddle-2 neurons could be independently modulated. Indeed, this is in line with model 1 and 3 predictions where Griddle-2 is modulated extrinsically (Fig. 3C,E, Fig. 4C, Extended Data Fig. 5E).

The converse experiments were also performed, i.e., inactivating Griddle-2 and imaging the activity in Handle-b (Fig. 4C4-C6). Handle-b responses to the mechanical stimulus were increased in normally fed larvae upon Griddle-2 inactivation (Fig. 4C5-C6, Extended Data Fig. 4F), which is consistent with Griddle-2 inhibition of Handle-b. In larvae fed on sucrose, the activity of the Handle-b neuron was not significantly different upon Griddle-2 inactivation and was similar to the activity upon Griddle-2 inactivation in larvae fed on standard food (Fig. 4C5-C6, Extended Data Fig. 4F). This lack of increase in Handle-b activity upon Griddle-2 inactivation in larvae fed on sucrose could be due to the different state of the network in the larvae fed only sucrose: the activity of Griddle-2 is already low in these larvae (Fig. 4A, Extended Data Fig. 4C), and removing it may not impact its inhibition of Handle-b. Moreover, when silencing the LNa type Griddle-2, other two LNa type neurons remain intact and may continue to inhibit Handle-b. In addition, removing Griddle-2 in this condition could impact the disinhibition of Basin-1 and/or the breaking of the positive feedback Basin-2-Handle-b-Basin-2.

### The feeding state-dependent increase in Basin-2 responses depends on the changes in activity of the inhibitory interneurons

Our previous work has shown that Basin-2 (B2) neurons receive input directly from the inhibitory neuron LNa and are disinhibited by the feedback inhibitory neuron Handle-b that makes inhibitory connections onto LNa and LNb neurons; Handle-b receives input primarily from Basin-2 neurons. The feedback disinhibition of Basin-2 creates positive feedback that stabilizes the Basin-2 ON state^35^. To determine whether the increase in Basin-2 responses in larvae fed on sucrose depends on feeding state-dependent changes in the activity of the inhibitory neurons, Handle-b neurons were optogenetically activated, and the activity of B2 neurons was recorded in response to a mechanical stimulus. We found that activating Handle-b in normally fed larvae increases the levels of Basin-2 responses to a level similar to that observed in larvae fed on sucrose (Fig. 5B, Extended Data Fig. 6B). We further silenced Handle-b neurons and monitored calcium responses in Basin-2 neurons in different feeding states (Fig. 5A, Extended Data Fig. 6A). Silencing Handle-b neurons in larvae fed on sucrose decreased Basin-2 responses and erased the difference in Basin-2 response levels between larvae fed on standard food and on sucrose (Fig. 5A, Extended Data Fig. 6A). This result is consistent with previous characterization of the Handle-b disinhibition of Basin-2. Because Handle-b disinhibits Basin-2, silencing Handle-b therefore increases the inhibition of Basin-2 by LNa, and thus reduces Basin-2’s responses to mechanical stimuli. In addition, in normally fed larvae where the activity of LNa is high, Basin-2 responses were almost completely abolished (Extended Data Fig. 6A).

**Fig. 5.**
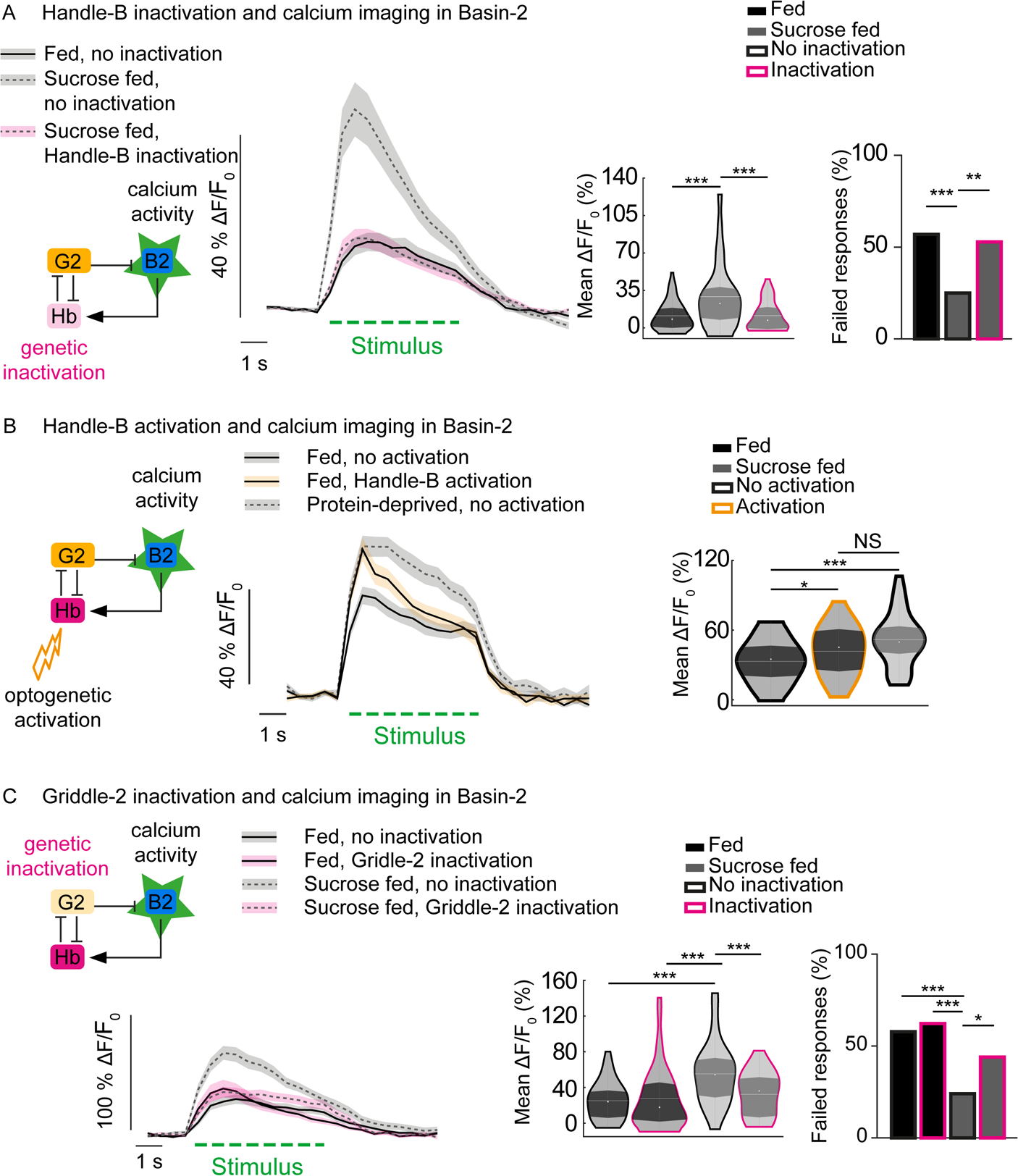
The modulation of inhibitory interneurons is required for the feeding state dependent changes in the circuit output. A: Left panel: mean calcium responses of Basin-2 over time from individuals fed on different diets, with (38H09-LexA>LexAop-GCaMP6s 22E09-Gal4>UAS-TNT) or without (38H09-LexA>LexAop-GCaMP6s +/>UAS-TNT) Handle-b inactivation. The green dashed line corresponds to stimulus onset. Middle panel: calcium response of Basin-2 averaged during the stimulus. White line represents the mean, white dot represents the median. Right panel: percentage of trials that failed to elicit a calcium response in Basin-2. Inactivating Handle-b in larvae fed on sucrose only prevents the effect of sucrose feeding on Basin-2 activity (n = 12-14 larvae, 5 trials per larva; ANOVA with post-hoc tests: ***: p < 0.001; chi-2 test ***: p < 0.001, **: p < 0.01). B: Left panel: mean calcium responses of Basin-2 over time from individuals fed on different diets, with or without optogenetic activation of Handle-b (22E09-Gal4>UAS-CsChrimson::tdTomato 38H09-LexA>LexAop-GCaMP6s) during the first second of mechanical stimulus (red box). The green dashed line corresponds to stimulus onset. Right panel: calcium response of Basin-2 averaged during the first second of the stimulus. White line represents the mean, white dot represents the median. Activating Handle-b in fed larvae phenocopies the effect of sucrose feeding on Basin-2 activity (n = 8 larvae, 5 trials per larva; ANOVA with post-hoc tests: ***: p < 0.001, *: p < 0.05). C: Left panel: mean calcium responses of Basin-2 over time from individuals fed on different diets, with (38H09-LexA>LexAop-GCaMP6s 55C05-Gal4>UAS-TNT) or without (38H09-LexA>LexAop-GCaMP6s +/>UAS-TNT) Griddle-2 inhibition. The green dashed line corresponds to stimulus onset. Middle panel: calcium response of Basin-2 averaged during the stimulus. White line represents the mean, white dot represents the median (ANOVA with post-hoc tests: ***: p < 0.001). Right panel: percentage of trials that failed to elicit a calcium response in Basin-2 (chi-2 test ***: p < 0.001, *: p < 0.05). See also Extended Data Fig. 6 and Supplementary Table 6.

Griddle-2 neurons were then inactivated and the activity of Basin-2 was imaged in the two feeding states. As expected, silencing Griddle-2 abolished the difference in Basin-2 responses in the two states. This result is consistent with the effect of Griddle-2 on Handle-b activity (Fig. 5). Silencing Griddle-2 in fed larvae resulted in small (but not significant) increase in Basin-2 responses (Fig. 5C, Extended Data Fig. 6C). However, the mean responses of Basin-2 in larvae fed only with sucrose unexpectedly and significantly decreased upon Griddle-2 silencing. We also computed Basin-2 response probabilities. While inactivating Griddle-2 did not affect Basin-2 response probabilities in normally fed larvae, it did so in larvae fed on sucrose; the non-response probabilities are significantly increased upon Griddle-2 inactivation in sucrose-fed larvae. (Fig. 5C, Extended Data Fig. 6C). The mean responses of trials with only Basin-2 ON responses reveal that the level of responses of Basin-2 was increased in normally fed larvae to the level found in larvae fed on sucrose in the control. The mean responses of trials with only Basin-2 ON responses also show that the level of Basin-2 response was decreased in larvae fed on sucrose (Extended Data Fig. 6C) and similar to the level of activity in fed control larvae.

These findings suggest that there are two mechanisms that control Basin-2 response probabilities and amplitudes, respectively. The two mechanisms are decoupled upon Griddle-2 inactivation. Inactivating Griddle-2 may decrease Basin-2 response probabilities by decreasing Basin-1 responses (that then disinhibits less Basin-2). At the same time, inactivating Griddle-2 increases Handle-b responses and thus amplifies Basin-2 responses more once Basin-2 responses are triggered.

Griddle-2 is thus required for state-dependent modulation of Basin-2, possibly by mediating the disinhibition of Basin-2 by Handle-b. Inactivating Griddle-2 in normally fed larvae disinhibits Basin-2. On the other hand, inactivating Griddle-2 in larvae fed on sucrose when the activity of Griddle-2 is already low (and Griddle-2 is not inhibiting Basin-2 strongly) may thus favor other inhibitory pathways and result in lower Basin-2 responses.

These results show that two types of reciprocally connected inhibitory neurons that promote competing actions inhibit each other during responses to a mechanosensory cue and are differentially modulated by the change in feeding states (starvation and feeding on sucrose). The activity of the Griddle-2 that promotes Hunching is decreased, while the activity of Handle-b that promotes Bending is increased. This differential modulation of the inhibitory neurons, in turn, has an impact on the state of the network in a feeding state-dependent manner and biases the behavioral responses towards less Hunching and more Head Casting by increasing the activity of the Basin-2 neurons when larvae are fed on sucrose only. In addition, it reveals that two different layers of the network are modulated by the changes of the feeding state: the reciprocally connected feedforward inhibitory neurons, as well as the feedback inhibitory neurons that stabilize the state of the network, resulting in the co-activation of both Basin-1 and Basin-2.

### The feedback inhibitory neuron receives input from an NPF-releasing neuron, that senses the changes in the feeding state

In order to determine the sources of feeding state-dependent modulation, we looked in the connectome for upstream partners of Griddle-2 and Handle-b neurons. Previous work has shown that the different neurons in the circuit receive input from long-range projection neurons that, as suggested by their morphology and connectivity, could bias the output of the network based on contextual/internal state information^35^. Interestingly, by matching light microscopy images to electron microscopy reconstruction images, we found one of these neurons is an NPF-expressing neuron that synapses on Handle-b. NPF (neuropeptide F), a homolog of the mammalian neuropeptide Y^46^, is a hunger signal, and its expression promotes feeding in both adult and larval Drosophila^27^. We, therefore, sought to investigate the implication of NPF neurons in modulating the activity of the inhibitory neurons in a state-dependent manner. In the larva, there are two pairs of NPF-expressing neurons. Both of them have cell bodies in the brain; the dorsolateral pair (DL-NPF) projects ipsilaterally within the brain lobes and the dorsomedial pair (DM) sends descending projections via the suboesophageal zone (SEZ) through the ventral nerve cord (VNC) (Extended Data Fig. 7A).

To study the influence of NPF neurons on Handle-b activity, the activity of the DM-NPF descending neuron was monitored in the different feeding states. To this end, we drove GCAMP6s expression in the NPF descending neurons with NPF-GAL4 and measured its intensity of fluorescence in the VNC projections of larvae fed either on standard food (fed), on 20% sucrose (sucrose fed) or completely starved (given only water for 90 minutes). We showed that starvation or feeding on sucrose both increase the activity of the NPF descending neuron (Fig. 6B).

**Fig. 6.**
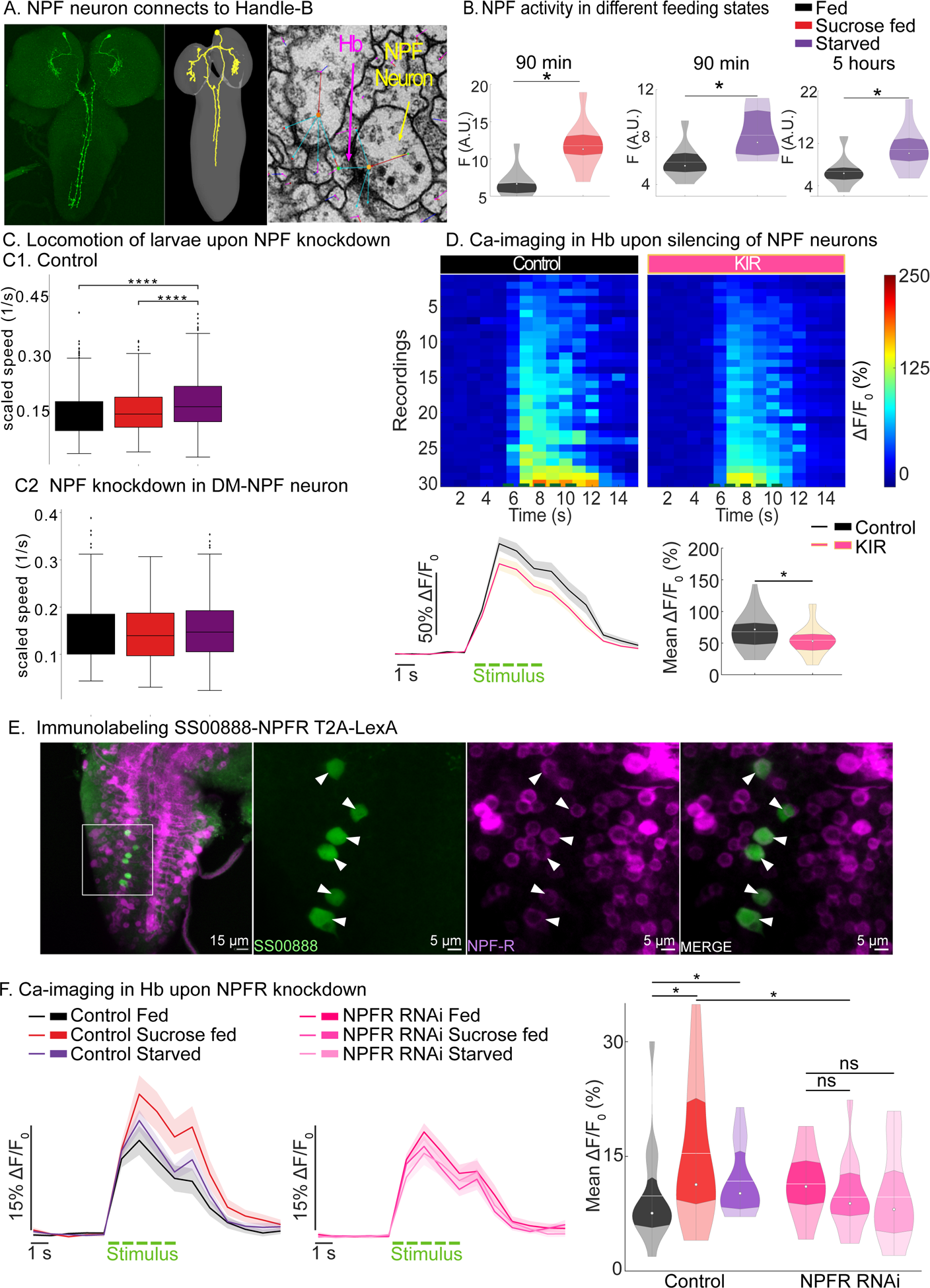
NPF-releasing neurons convey internal state information to the feedback inhibitory neuron. A: Expression profile characterization of GMR_SS01635>UAS-GFP stained with a GFP-specific antibody. GMR_SS01635 split-Gal4 line selectively targets one pair of NPF neurons. Comparison of light microscopy images with electron microscopy (EM) reconstruction images indicates that a descending neuron synapses on Handle-b (Jovanic *et al*, 2016) is the DM-NPF descending neuron. EM image shows neuropeptide-containing dense core vesicles in the NPF descending neuron near one of its synapses with Handle-b. B: Baseline calcium fluorescence measured in the projections of NPF descending neurons in the VNC (ventral nerve cord) of larvae fed on sucrose only for 90 minutes (N = 10 larvae), starved for 90 minutes (N = 10 larvae) or 5 hours (N = 8-9 larvae). White line represents the mean, white dot represents the median. Starvation or feeding only on sucrose only increase NPF descending neuron activity (two-tailed T-test: *: p < 0.05). C: Analysis of larval locomotion in the absence of sensory stimulus. Top row shows control larvae (C1). Bottom row shows larvae with NPF knockdown in DM-NPF descending neurons (C2). NPF knockdown in DM-NPF descending neuron abolishes differences in locomotion mean between larvae in different states; as measured by the average movement speed (Mann-Whitney test, ****: p<0.0001, ***: p < 0.001, **: p < 0.01, *: p < 0.05) D: Calcium responses in Handle-b with (UAS-GCamP6s; 22E09-Gal4, NPF-LexA>LexAop-KIR) or without (UAS-GCamP6s; 22E09-Gal4, NPF-LexA;+) NPF neurons silencing. Left panel: calcium responses of Handle-b, in each trial of mechanosensory stimulation. Right lower panel: mean calcium response of Handle-b over time. The green dashed line corresponds to the stimulus. Top right panel: mean calcium response averaged during the stimulus. White line represents the mean, white dot represents the median. Stimulus-induced activity of Handle-b is decreased upon Coaster silencing (n = 10 larvae, 3 trials per larva, t-test p = 0.045). E: Immunohistochemical labeling for NPFR in Handle_B UAS-GCamP6s is expressed in Handle-b using the GMR_SS00888 split-Gal4 line (green) and LexAop-jRGeco1a is expressed under the control of the NPFR promoter using a T2A-LexA construct (magenta). Antibodies against GFP and dsRed were used to increase detection sensitivity. Co-localization of LexAop-jRGeco1a and GCamP6s show that Handle-b expresses the NPFR. F: Calcium responses in Handle-b upon NPFR knockdown (GMR_SS00888>UAS-NPFR-RNAi; UAS-GCamP6s) compared to a control (GMR_SS00888>UAS-GCamP6s). Left panel: mean calcium response of Handle-b over time in control larvae fed, fed on sucrose or starved. The green dashed line corresponds to the stimulus. Middle panel: mean calcium response of Handle-b over time upon NPFR knockdown in larvae fed, fed on sucrose or starved. The green dashed line corresponds to the stimulus. Right panel: mean calcium response averaged during the stimulus. White line represents the mean, white dot represents the median (n = 14-15 larvae). Handle-B responses are increased in control larvae upon feeding on sucrose (Mann-Whitney test, p = 0.04846) or starvation (Mann-Whitney test, p = 0.02807). NPFR knockdown erases this modulation (Mann-Whitney test, p = 0.09853 and p = 0.121).NPFR knockdown significantly decreased Handle-B responses in larvae fed on sucrose (Mann-Whitney test, p = 0.03754). See also Extended Data Fig. 7 and Extended Data Fig. 8 and Supplementary Tables 2, 3 and 6.

NPF was shown to be involved in promoting feeding^47^. To investigate whether the increase in activity of NPF descending neurons upon 90-minute sucrose feeding or starvation had an influence on NPF release and behavior, NPF was knockdown selectively in the NPF descending neuron using a split-GAL4 driver that labels only the descending pair of NPF neurons (Fig. 6A) and larval locomotion monitored in larvae fed on standard food, larvae fed on sucrose and starved larvae (Fig. 6C, Extended Data Fig. 8A). Downregulating NPF in the DM-NPF neurons descending into the VNC impaired the increase in exploration observed in larvae fed on sucrose and starved larvae: it abolished the difference in tortuosity over 20 s, time fraction in runs, mean scaled speed and maximum dispersal over 60 s between larvae fed on standard food and starved larvae. Moreover, NPF downregulation abolished the difference between larvae fed on standard food and larvae fed on sucrose or even decreased exploration in larvae fed on sucrose compared to larvae fed on standard food as observed by increased tortuosity over 20 s and decreased maximum dispersal and time fraction in runs compared to larvae fed on standard food (Fig. 6C, Extended Data Fig. 8A).

To understand the influence of NPF neurons on Handle-b activity, the interactions between the two neurons were tested functionally by hyperpolarizing the NPF neurons with the inwardly rectifying potassium channel Kir2.1 using an NPF-LexA driver line that labels both pairs of NPF neurons and monitoring calcium responses of Handle-b to a mechanical stimulation. Upon NPF neurons silencing, we observed a decrease in Handle-b responses to mechanical stimulations (Fig. 6D). The NPF descending neuron could thus facilitate responses of Handle-b neurons. Upon changes in the feeding state, the NPF neurons whose activity increases upon starvation and sucrose feeding, could enhance Handle-b responses to a mechanical stimulus, resulting in higher Handle-b responses (Fig. 6B, D) that bias behavioral choice towards less Hunching and more Head Casting.

In the connectome, we found large dense core vesicles in the NPF neurons in the proximity of the Handle-b neuron (Fig. 6A). Moreover, we showed that Handle-b expresses the NPFR1 using a T2A-LexA method^48^ where we drove the expression of one reporter in all NPFR1 neurons using the NPFR1 T2A-Lexa and of another reporter with a Handle-b specific split-GAL4 driver. The intersection of the two expression patterns revealed that the Handle-b neurons express the NPFR1 (Fig. 6E). Having established that Handle-b and Griddle-2 are instrumental in the feeding state-dependent modulation of the behavioral choice of larvae in response to mechanical stimulation and that NPF modulates Handle-b, given that the model and functional connectivity experiment predict that the Handle-b and Griddle-2 could be independently modulated (Fig. 4C, Extended Data Fig. 5E-F), we investigated whether Griddle-2 could be also influenced by NPF. We found that Griddle-2 does not receive synaptic input from the NPF descending neurons. Since neuropeptides can act outside of synaptic sites and we found dense core vesicles characteristic for neuropeptide release along the NPF axon in the proximity of Griddle-2 neurons, we also examined the NPFR1 expression in Griddle-2 neurons, as well as in Basin-1 and Basin-2 using a similar approach as for Handle-b and didn’t find any NPFR1 positive cells in these neurons (Extended Data Fig. 7D).

We then downregulated the expression of NPFR1 in the Handle-b neurons and imaged their calcium responses to mechanical stimulation in larvae fed on standard food, larvae fed only on sucrose and in starved larvae (Fig. 6F, Extended Data Fig. 8B). Downregulation of NPFR1 in Handle-B abolished the feeding-state dependent modulation of Handle-b as there was no significant increase in Handle-B responses upon sucrose feeding or starvation when NPFR1 was downregulated (Fig. 6F, Extended Data Fig. 8B).

Altogether, these results show that NPF neurons sense the changes in feeding state and their activity increases upon starvation or sucrose only feeding. The changes in NPF activity influence Handle-b activity that increases upon starvation or sucrose only feeding due to a facilitating effect of NPF. This increase in activity then biases the sensorimotor decisions towards less Hunching and more Head Casting.

### sNPFR in the interconnected inhibitory interneurons mediates the state-dependent modulation of larval behavioral responses to a mechanical stimulus

The model and calcium imaging suggests that Griddle-2 and Handle-b are independently modulated by the feeding state (Fig. 4). Since Griddle-2 does not express the NPFR1 receptor, other neuropeptides could influence Griddle-2 and the circuit. We therefore investigated the effect of another neuropeptide on the circuit: sNPF (the short neuropeptide F). sNPF is a second homolog of the mammalian NPY whose receptor is widely distributed in the larval nervous system, including the VNC^49^, and was shown to be involved in regulating hunger-driven behaviors^27^, and in facilitating mechano-nociceptive responses^37^ among other functions. Therefore, we sought to examine whether the interneurons in the circuits express the receptor for sNPF. By genetically co-expressing a red fluorescent reporter (jRGeco1a) under the control of LexA drivers that label either Handle-b, Griddle-2, Basin-1 or Basin-2 neurons we identified in the literature and existing Gal4 databases (see Methods for details), and a green fluorescent reporter (GCamP6s) under the control of the sNPFR1 promoter with a T2A-Gal4 construct, we showed that both Handle-b (Fig. 7A) and Griddle-2 (Fig. 7D) express the sNPFR1, while the two Basin neurons do not (Extended Data Fig. 7E).

**Fig. 7.**
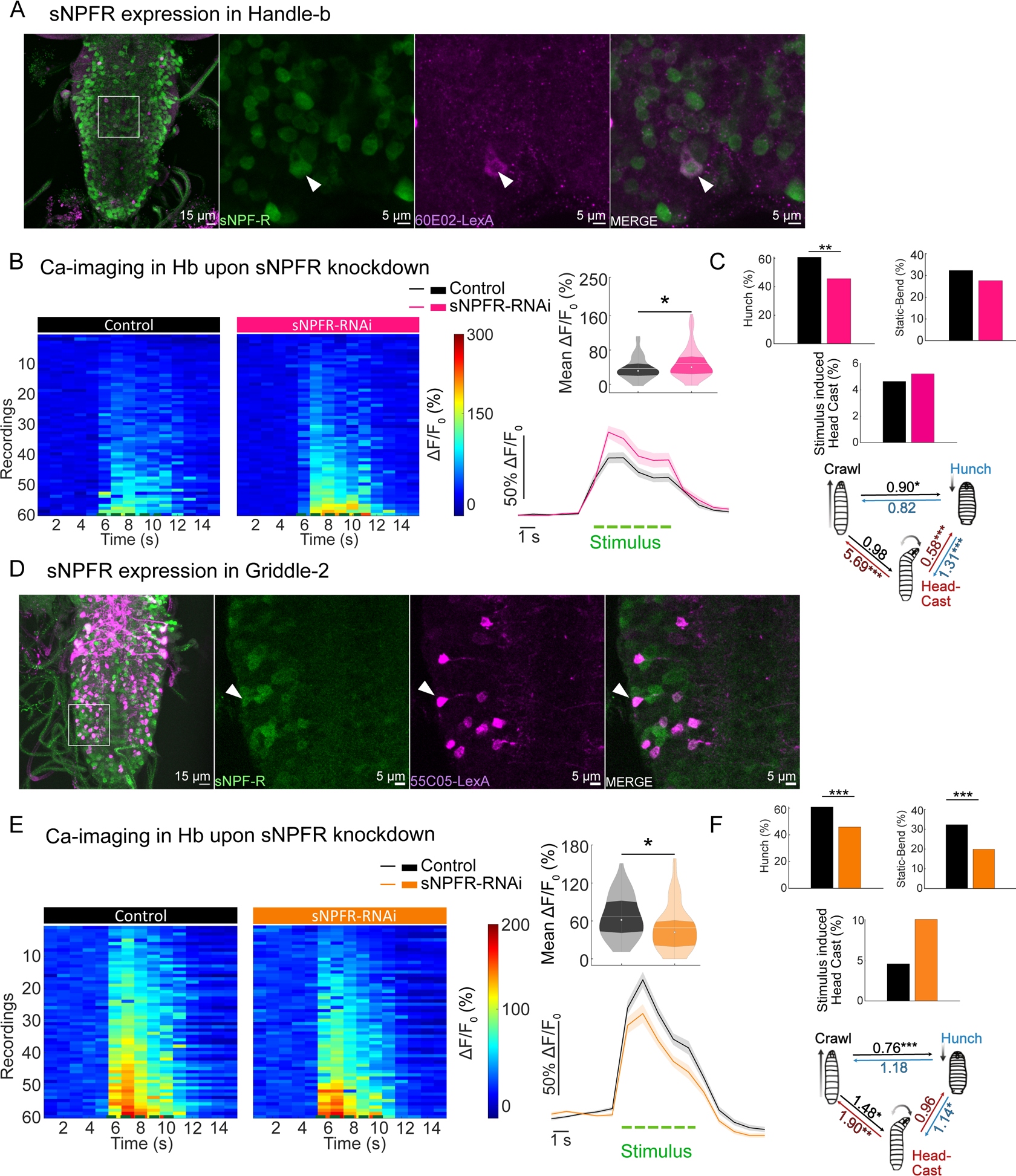
sNPF signaling inhibits Handle-b and facilitates Griddle-2 mechanosensory responses. A: Immunohistochemical labeling for sNPFR in Handle-b. LexAop-jRGeco1a is expressed in Handle-b using the 60E02-LexA line (magenta) and UAS-GCamP6s is expressed under the control of the sNPFR promoter using a T2A-Gal4 construct (green). Antibodies against GFP and dsRed were used to increase detection sensitivity. Co-localization of antibodies against jRGeco1a and GCamP6s shows that Handle-b expresses the sNPFR. B: Calcium responses in Handle-b upon sNPFR knockdown (GMR_SS00888>UAS-GCamP6s; UAS-sNPFR-RNAi) compared to a control (GMR_SS00888>UAS-GCamP6s) in Handle-b. Left panel: calcium responses of Handle-b from different individuals and trials. One line is a trial. Lower right panel: mean calcium response of Handle-b over time. The green dashed line corresponds to the stimulus Top right panel: mean calcium response averaged during the stimulus. White line represents the mean, white dot represents the median. Stimulus-induced activity of Handle-b is increased by sNPFR knockdown (n = 15 larvae, 4 trials per larva, t-test p = 0.035). C: Behavior in response to air-puff during the first five seconds upon stimulus onset,for larvae in which sNPFR was knocked down in Handle-b neurons (SS00888>sNPFR-RNAi) compared to the control (n = 185-260 larvae) (***: p < 0.001, **: p < 0.01, *: p < 0.05). D: Griddle-2 immunostaining for sNPFR expression. LexAop-jRGeco1a is expressed in Griddle-2 using the 55C05-LexA line (magenta) and UAS-GCamP6s is expressed s expressed under the control of the sNPFR promoter using a T2A-Gal4 construct (green). Antibodies against GFP and dsRed were used to increase detection sensitivity. Co-localization of antibodies against jRGeco1a and GCamP6s show that Griddle-2 expresses sNPFR. E: Calcium responses in Griddle-2 upon sNPFR knockdown (SS_TJ001>UAS-GCamP6s; UAS-sNPFR-RNAi) compared to the control (SS_TJ001>UAS-GCamP6s). Left panel: calcium responses of Griddle-2 from different individuals and trials. One line is a trial. Lower right panel: mean calcium response of Griddle-2 over time. The green dashed line corresponds to the stimulus. Top right panel. mean calcium response averaged during the stimulus. White line represents the mean, white dot represents the median. Stimulus-induced activity of Griddle-2 is decreased by sNPFR knockdown (n = 10 larvae, 3 trials per larva, t-test p = 0.003). F: Behavior in response to air-puff during the first five seconds upon stimulus onset, for larvae in which sNPFR was knocked down in Griddle-2 neurons (SS_TJ001>sNPFR-RNAi) compared to the control (n = 243-438 larvae, ***: p < 0.001, **: p < 0.01, *: p < 0.05). See also Extended Data Fig. 7, Extended Data Fig. 9 and Supplementary Tables 4, 5 and 6.

To determine whether sNPF signaling was involved in responses to a mechanical stimulus in these two neurons, their responses were monitored upon sNPFR1 downregulation. Calcium imaging recordings show that genetically downregulating sNPFR1 expression in Handle-b increases its responses to mechanical stimulation (Fig. 7B), suggesting that sNPFR1 has an inhibitory effect on Handle-b. On the other hand, responses of Griddle-2 were decreased upon sNPFR1 downregulation (Fig. 7E), suggesting that sNPF facilitates Griddle-2 responses to a mechanical stimulus. Since Handle-b inhibits Hunching and promotes Head Casting, while Griddle-2 promotes Hunching and inhibits Head Casting, downregulating the sNPFR1 in these neurons would inhibit Hunching and promote Head Casting. Indeed, the behavior experiments showed that sNPFR1 knockdown in either of these two neurons leads to less Hunching and more transitions to Head Casting (Fig. 7C, F) in response to an air-puff. The sNPFR1 knockdown in Handle-b and Griddle-2 leads to a modulation of their activity similar to that caused by feeding on only sucrose (lower Griddle-2 and higher Handle-b activity) and yields the same behavioral outcome. This suggests that sNPF signaling targeting these neurons could be downregulated in larvae fed on sucrose and at least partially responsible for the modulation of the behavioral choice of the larvae in response to a mechanical stimulation. Giving stock to this hypothesis, the downregulation of sNPFR1 in Handle-b or Griddle-2 did not impact their calcium responses to mechanical stimulation in larvae fed on sucrose or starved (Extended Data Fig. 9).

Altogether these results show that Handle-b and Griddle-2, reciprocally interconnected inhibitory neurons that drive opposing actions, are differentially modulated by the sNPF signaling pathway to bias the response to air-puff towards less Hunching and more Head Casting.

## Discussion

Whereas many studies showed that various behaviors are affected by an animal’s physiological state, especially hunger, this work goes further by describing the neural circuit mechanisms by which transient changes in the physiological status impact behaviors that are not directly related to feeding itself. To determine whether and how changes in the feeding state affect behaviors unrelated to feeding, we took advantage of a well characterized behavioral response of *Drosophila* larva to a mechanical stimulus for which we had in the past dissected the circuit mechanisms underlying the selection between two main types of responses: Hunch and Head Cast. The current study reveals the neural circuit mechanisms of modulation of these sensorimotor decisions by the feeding conditions. Slight changes in feeding conditions affect an animal’s motivational state and bias responses to an air-puff towards less Hunching and more Head Casting. Reciprocally interconnected inhibitory neurons that drive competing actions are differentially modulated by the changes in the feeding state: the activity of the neuron that promotes a Hunch is decreased and the activity of the neuron that inhibits a Hunch is increased. This modulation at the level of inhibitory neurons involving NPF and sNPF signaling systems biases the output of the network towards promoting Head Casting and inhibiting Hunching, a modulation consistent with the state-dependent behavioral changes we observed.

Hungry animals behaving differently in non-feeding related contexts have been reported in various studies and across the animal kingdom^7,51^. However, the underlying logic and neural mechanisms have not been well understood. These changes could be linked to overall changes in behavioral strategies in hungry individuals^7^ to ensure survival. Risk-taking has been shown to be increased by hunger^50–52^, even in social-related decisions^7,53^, further raising questions about how food-deprivation signals are integrated into the neural computations underlying non-feeding related behaviors. Animals need to be able to make any decisions flexibly depending on the need that is most critical at a given moment. Our study shows that a circuit underlying sensorimotor decisions in response to a mechanical stimulus located at the early stages of sensory processing is influenced by the changes in diet. This finding supports the idea that state dependent flexibility of behavior could be achieved by the physiological state acting on circuits throughout the nervous system and thus reorganizing its activity in a distributed manner. In various ecological contexts, survival often entails a trade-off between avoiding dangers and pursuing food and water-seeking behaviors^51^. For example, increased exploration will increase the likelihood of finding food but also of encountering a dangerous situation^51^. If animals’ need for food or water increases either due to food deprivation or thirst, they might be more likely to explore intensively despite an increasing risk of threat. Similarly, food-deprived and thirsty animals might take more risks and ignore aversive cues to increase their chances of getting near food or water sources. Hungry animals use different strategies when escaping predators compared to satiated ones^17,23^.

While various studies have shown that hunger affects behaviors by altering the responses of sensory neurons^9,11,54,55^, others have implicated central mechanisms in feeding state dependent behavioral flexibility^13,31,56^. We found that the activity of chordotonal mechanosensory neurons that sense the air-puff was not significantly altered upon feeding on sucrose or starvation, contrary to the activity of the downstream neurons. The changes at the level of sensory pathways tunes animals’ perception in order to increase their likelihood of finding food and feeding by increasing responsiveness to appetitive and decreasing responsiveness to aversive food-related stimuli. The implication of central mechanisms, on the other hand, may suggest that hunger acts as a global regulator of behavior, i.e. hunger may change brain activity in such a way to reevaluate goals and behavioral strategies in order to increase animals’ chances of survival^7^.

Our experiment monitors calcium responses of all the chordotonal sensory neurons together. The different inhibitory and projection neurons in the circuit receive inputs from different chordotonal subtypes. This could also explain the differential modulation of the different neuron subtypes in the circuit if some subtypes of chordotonal are modulated by the changes in the internal state while others are not. However, optogenetic activation of all the chordotonal neurons in the different feeding conditions still resulted in decreased Hunching and increased Head Casting upon starvation or sucrose diet, strongly suggesting that downstream neurons are involved. Even if chordotonals are involved in altering behavior in a state-dependent manner, their modulation is not required to alter the behavior in response to the mechanical stimulus due to the contribution of the downstream circuitry.

Calcium imaging combined with neuronal manipulations revealed that, in the circuit for selecting between the Hunch and the Head Cast, inhibitory neurons (and not the projection neurons) are the target of modulation by the changes in the feeding conditions. Inhibitory neurons were shown to be the target of contextual modulation in other systems^57–59^.

Previous work has identified reciprocal inhibition of inhibition as a motif underlying the competition between a startle-type action and an exploratory action^35^. Such a motif was shown to underlie similar computations in different species and brain areas^35,60–65^ and was proposed to provide flexibility to the selection process^35,63,65^. The current study determines that one of the reciprocally connected inhibitory neurons within this motif, LNa type Griddle-2, is modulated by the changes in the feeding conditions and that this modulation contributes to biasing sensorimotor decisions. These results confirm theoretical predictions that such a motif confers the sensorimotor circuit the capacity to be tuned to other types of information, in this case, internal state information (starvation and thirst). Moreover, these findings are in line with predictions that shaping the output of the network through disinhibition by reciprocally connected inhibited neurons allows for flexible, competitive selection^35,63,65^.

This work shows that another type of inhibitory neuron, Handle-b, which is a feedback inhibitory neuron that provides positive feedback to the Hunch inhibiting Basin-2 neuron through feedback disinhibition, is also modulated by changes in internal physiology. Handle-b is also reciprocally connected to the LNa neurons that participate in the reciprocal inhibition of inhibition motifs. This connectivity pattern suggests that, in addition to competition within the reciprocal inhibition of inhibition motifs, the competition between the two layers of the circuits is also modulated by the changes in internal physiology. Similar to recurrent excitation, the feedback disinhibition motif provides positive feedback that stabilizes the selected output^35^. Using inhibitory rather than excitatory connection may have the advantage of allowing the decisions to be influenced by contextual and state information. In addition, in this circuit architecture, the feedback inhibitory neuron Handle-b contacts both reciprocally connected inhibitory neurons in the circuit and can thus shape the circuit activity at the level of the site of competition. It is thus well suited to be the site of integration of mechanosensory information and information about an animal’s state.

The behavioral response of the larvae to a mechanical cue depends on the state of the circuit at the level of reciprocally interconnected inhibitory neurons, which will shape the activity of the Basin projection neurons to either give rise to a state where Basin-1 only is active or a state where Basin-1 and -2 are co-active^35^. We showed that the feeding state-dependent modulation of the Basin-2 neuron is dependent on the modulation of the activity of Handle-b and Griddle-2. Accordingly, genetic co-labeling of Basin-1 or Basin-2 neurons with either sNPFR1 or NPFR1 using the T2A GAL4/LexA technology showed no expression of these neuropeptide receptors in the two Basin neurons. This is in line with the finding that the state-dependent changes in the air-puff induced sensorimotor decisions are caused by the modulation that acts on the reciprocally interconnected inhibitory neurons that integrate mechanosensory information with current internal state needs.

Previous work revealed that the inhibitory neurons in the mechanosensory circuit are contacted by long-range projection neurons^35^. These long-range projection neurons may carry contextual or state information to the mechanosensory circuit. We found that among these long-range projection neurons is a pair of NPF-releasing neurons that contacts the Handle-b. NPF is a homologue of the mammalian neuropeptide Y that is a hunger signal. NPF neurons have indeed been shown to be involved in hunger dependent behaviors in both adult flies and larvae^27,47^. We found that the activity of the NPF descending pair of neurons changes as a function of the feeding state. These neurons could thus convey the information about the satiation state to the mechanosensory circuit and bias the sensorimotor decisions to mechanosensory cues by modulating the activity of the Handle-b. Additionally, the release of NPF at the proximity of the circuit by the NPF neurons (as suggested by the existence of dense core vesicles) could modulate Handle-b that expresses the NPFR1 (while other neurons in the circuit do not). The fact that the feedback inhibitory neuron is directly influenced by the NPF neurons supports the idea that it could gate circuit activity in a state dependent manner. Our results indicate that both the feeding state induced changes in locomotor strategy (motivational locomotor state and exploration persistence) and the acute response to transient environmental stimuli are dependent on NPF signaling. The descending NPF neurons could convey information about the internal state to diverse neuronal populations in the VNC to regulate exploratory locomotion and stimulus-dependent motor responses, thus adjusting behavioral interactions with the environment according to the current animal needs. The NPF could serve as an internal state signal that couples various sensorimotor behaviors to the motivational/exploratory state of the animal and its physiological needs, thus regulating behavior across different timescales.

## Acknowledgments

### Funding

This work was supported by ANR PIA funding: ANR-20-IDEES-0002 (T.J), Agence Nationale de la Recherche: ANR-17-CE37-0019-01 (T.J.), ANR-NEUROMOD (ANR-22-CE37-0027) (T.J.), ANR-NEUC-0002-01 (T.J), Fédération pour la recherche sur le cerveau (FRC) (T.J.), Équipe FRM EQU202303016317 (TJ), Fondation des Treille (E.T.), D. M. received a PhD fellowship from the Paris-Saclay University; Tramway, ANR-17-CE23-0016 (J.B.M), the inception Project PIA/ANR-16-CONV-0005,OG (J.B.M), Investissement d’avenir programme under the management of ANR, ANR-19-P3iA-0001 (PRAIRIE 3IA Institute (J.B.M), German Federal Ministry of Education and Research (BMBF DrosoExpect, 01GQ2103A, M.N.), Ministry of Culture and Science of the State of Northrhine Westphalia (iBehave, Netzwerke 2021, M.N.). P.S. received a PhD stipendship from the German Research Foundation (DFG-RTG 1960, 233886668, M.N.).

The funders had no role in study design, data collection and analysis, decision to publish, or preparation of the manuscript.

### Author contributions

E.T. behavioral and physiology experiments and analysis, Calcium-imaging experiments and analysis; Writing: figures, edits and revisions; D. M. behavioral experiments and analysis, Calcium-imaging experiments and analysis, immunohistochemistry experiments, Writing: original draft, figures; A.B. modeling; P.S. Data analysis; C.B. behavioral classification and analysis, statistical analysis, writing, methods; F. V. behavioral experiments; V. S. behavioral and physiology experiments; A.H. and S. A immunohistochemistry experiments; M.N. supervision, funding acquisition; J.B. Methodology, supervision, funding acquisition, writing-edits and revision; T.J. Conceptualization, analysis, supervision, funding acquisition and project administration; writing: original draft, figures, edits and revision.

## Material and methods Drosophila rearing and handling

Flies (*Drosophila Melanogaster*) were raised on a standard food medium (ethanol 2%, methylhydroxybenzoate 0.4%, yeast 8%, cornmeal 8%, and agar 1%) at 18°C. Third instar larvae were collected as follows: male and female flies from the appropriate genotypes were placed together for mating, then transferred at 25°C for 12-16 h on a petri dish containing a fresh food medium for egg laying. The petri dish was then placed at 25°C for 72 h. Foraging third instar larvae were collected from the food medium by using a denser solution of 20% sucrose, scooped with a paint brush into a sieve and gently and quickly washed with water. Larvae used for optogenetic experiments were raised at 25°C in complete darkness, on standard food supplemented with all-trans retinal at 0.25 mM (R240000, Toronto Research Chemicals). The full list of genotypes used in the study can be found in the Supplementary Table 1 Resource table.

## Dietary treatments

For dietary treatments, larvae were placed in 60×15 mm circular petri dishes that contained a 45 mm circular Whatman paper. Larvae were subjected to different diets: standard food without agar for “fed” larvae (as described in the Drosophila rearing section), 20% sucrose solution for “protein deprived” larvae, and water for “starved” larvae. The Whatman paper was soaked with 0.6 mL MilliQ water (“starved”), sucrose solution (“fed on sucrose”), or soaked with 0.6 mL water and 1 mL of standard food medium was added on top (“fed”). Larvae were collected after the appropriate amount of time (90 or 300 min) and rinsed in water before behavioral, imaging, or biochemistry experiments. For the behavioral experiments with rehydration, larvae were collected after 90 minutes, rinsed in water, and placed in a petri dish with a Whatman paper soaked with 0.6 mL water for 15 minutes. Likewise, for the refeeding experiments, starved larvae were collected after 300 minutes, rinsed, and placed for 15 minutes in a petri dish with standard food medium and water as in the “fed” condition. After the treatment, larvae were once more collected and rinsed before the experiment.

## Behavioral tracking

We used an apparatus previously described^44,45^. Briefly, the apparatus comprises a video camera (Basler ace acA2040-90 µm) for monitoring larvae, a ring light illuminator (Cree C503B-RCS-CW0Z0AA1 at 624 nm in the red), a computer and a hardware module for controlling air-puff, controlled through multi worm tracker (MWT) software (http://sourceforge.net/projects/mwt)^45,66^. The arena consisted of a 25625 cm2 square of 3% Bacto agar gel (CONDALAB 1804-5) with charcoal (Herboristerie Moderne, 66000 Perpignan) in a plastic dish, and was changed for each experiment. For optogenetic experiments, plates without charcoal were used, and larvae were tracked thanks to IR light. Collected third instar larvae were washed with water, moderately dried and spread on the agar starting from the center of the arena. We tested approximately 30–100 larvae at once during each experiment. The temperature of the behavioral room was kept at 25°C.

## Locomotion analysis

To assess larval locomotion in the absence of stimulation, larvae were placed on top of the agar in the arena inside the tracker, and either tracked for 5 min continuously (intact attP2>UAS-TNT larvaewild type larvae) or for 60s. The coordinates of the 11 points along the central spine and the outline of each larva were computed as described previously^44,45^. These 2D X-Y coordinates, structured as time series of irregular framerate, labeled by the instantaneous tracking time of the recording, comprise the raw datasets which have been subsequently analyzed to derive all secondary metrics.

Analysis was performed in python using the *<u>larvaworld</u>* behavioral analysis and simulation platform (https://pypi.org/project/larvaworld/). The 3-step analysis pipeline included preprocessing, computation of secondary angular, translational and temporal metrics and behavioral epoch detection to annotate strides, runs and pauses, as described previously^67^. During preprocessing the raw time series were adjusted to a 10 Hz constant framerate by interpolating them at a 0.1 second timestep. Noise reduction was achieved by applying a low-pass filter with a 2Hz cut-off frequency, a threshold high enough not to alter the crawling-related dynamics around the dominant ∼1.4Hz crawling frequency.

For trajectory-based spatial metrics such as pathlength and dispersal, to avoid the cumulative effect of body micromovements, the position of the 9^th^ point along the midline was used as a proxy for the larva’s position, a relatively stable rear point unaffected by lateral and translational jitter. To correct for different larval sizes, any metric measured in absolute spatial units (m or mm) can be scaled to body-length, measured in dimensionless body-length units. As the instantaneous body-length of an individual larva fluctuates during crawling due to subsequent stretching and contraction, individual larva length is defined as the median of the midline length across time (total length of the line connecting all 11 midline points). A trajectory’s pathlength is the cumulative displacement of the larva during the entire track. Dispersal is the instantaneous straight-line distance relative to its initial position. Track tortuosity was quantified by the straightness index (S.I), computed by advancing a fixed time window (20 seconds in this study) along the track and calculating at each point the ratio of the dispersal to the actual distance traveled. This index, which varies from 0 (no movement) to 1 (straight line movement), can capture very well the complexity of the movement at various scales (set by the window time frame) throughout the track.

For the detection of peristaltic strides and crawl-pauses, the scaled crawling speed time series were used. To this end the dominant crawling frequency for each track was extracted by applying a Fourier analysis and its inverse was used as the expected duration of a peristaltic cycle. A stride was therefore defined as the epoch between two local speed minima, that included a local maximum of at least 0.3 body-lengths/s and lasted between 0.7 and 1.5 times the expected cycle duration. A run was defined as an uninterrupted sequence of consecutive strides and a crawl-pause as an epoch lacking any strides during which the scaled speed was constantly below the 0.3 body-lengths/s threshold.

## Air-puff stimulation during behavioral tracking

air-puff was delivered as described previously^35,44,45^ to the 25625 cm2 arena at a pressure of 1.1 MPa through a 3D-printed flare nozzle placed above the arena, with a 16 cm x 0.17 cm opening, connected through a tubing system to plant supplied compressed air. The strength of the airflow was controlled through a regulator downstream from the air amplifier and turned on and off with a solenoid valve (Parker Skinner 71215SN2GN00). Air-flow rates at 9 different positions in the arena were measured with a hot-wire anemometer to ensure consistent coverage of the arena across experimental days. The air-current relay was triggered through TTL pulses delivered by a Measurement Computing PCI-CTR05 5-channel, counter/timer board at the direction of the MWT. The onset and duration of the stimulus were also controlled through the MWT. Larvae were left to crawl freely on the agar plate for 60 seconds prior to stimulus delivery. air-puff was delivered at the 60th second and applied for 30 seconds.

## Behavior classification and analysis

Behaviors were detected thanks to a custom-made machine learning algorithm that was previously described^44^. Behaviors were defined as mutually exclusive actions. Larvae were tracked using MWT software, all the time series of the contours and the spine of individual larvae are obtained using Choreography. From these times series some features are computed, center of the larva, velocities, etc., all key features are presented in Masson *et al.*, 2020. Behavior classification consists of a hierarchical procedure that were trained separately based on a limited amount of manually annotated data. Here, we required a more detailed definition of behavior. Hence, we extended the hierarchy with another layer to separate some Bends and Hunches, between different behaviors. Bends were separated into Static Bend, Head-Cast (see the description of each behavior below). We take all the Bends, Hunches and Back-ups obtained by the first classification algorithm, to reclassify with new annotated data on new lines.

### Action definition

#### Head Cast

dynamic bends in which the head moves laterally from one side to the other. There are two exits from Head-Cast in both the head moves strongly, but one that is slower and the tail moves at the same speed as the center of mass, and on the contrary a second one where the tail moves a lot (fast Head-Cast).

#### Static Bend

Low speed turning movement, and where the head moves little, and the angle between the segment between the center of mass and the head and the center of mass and the tail remains constant.

### New annotated data

We required new annotated data to ensure the classifier matched the phenotype of the larva used in these experiments. The sets of, Hunch, Bends, Backs and Crawls were manually tagged from actions selected using the Masson *et al,* 2020 old behavior classification pipeline. A few numbers of tags are used for the model.

### New features

To train the new layer of the pipeline, we combined features that were previously evaluated in the classifier’s preceding layer with new features. Each characteristic is calculated for the time step we are examining and the three time steps before and after.

- The three velocities include the head velocity, the motion velocity, and the tail velocity, all normalized by the length of the larva. The motion velocity is the velocity of center of mass return by the MWT software. Head and tail are the terminal point of the spine; The averaging along the spine curve and its derivative, 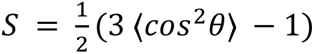 with *cos θ* the scalar product between normalized vectors associated to a segment of the spine and the direction of the larva body.
- The shape factor 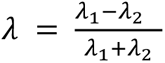 with *λ_i_* the eigenvalues of the mean covariance matrix of movement which characterizes the shape of the larva and takes value between 0 and 1.

We have also introduced new features:

- The ratio between the length of the head-center of mass and the tail-center of mass 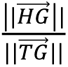 with *H,T* and *G* respectively coordinate points of the head, the tail and the center of the masse
- The projection of the head and tail velocity on the spine of the larva. If we note the velocity vector of the head *HV_h_* with H coordinate point of the head and *V_h_* the coordinate point at the end of the velocity vector, the projection point satisfies the basic relationship: 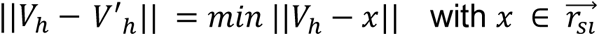.
- The cosine of the angle between the vector of the head (tail) velocity and the first (last) segment of the larva. 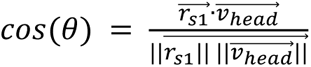 with 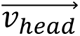 the vector velocity of the head and 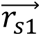 the first vector of the spine.

All features are normalized by the length of the larva to ensure scale-free properties.

### New classifications

We employed a Random Forest algorithm for classification, distinguishing each time step of original Bends, Hunches, and Backs into new categories. We conducted ten random forests on all Hunch and back tags, along with a random selection of a thousand bends, utilizing balanced weights. The predicted behavior represents the most probable outcome, with each random forest’s confusion matrix exceeding 80% accuracy for each behavior.

To address the issue of having a behavior span only two time steps, which is not biologically plausible, we introduced a preventive measure by adding five time steps before and after the behavior. Additionally, to mitigate noisy results, we implemented smoothing. This involved excluding behaviors with fewer than three time steps, logically categorizing them as the behavior before or after (based on the length of the behavior and behaviors N+2 and N-2). According to our knowledge, Hunch behavior initiates at the beginning of stimulation, around 60 second^35,44^.

Larvae typically do not exhibit multiple Hunches. In cases where they do, we classify the second Hunch as a cast, a classification verified through ground truthing. We applied the same threshold as outlined in Masson *et al.,* 2020 to the effective length change during the behavior. If a Hunch fails to surpass this threshold, the time window is assigned to the small behavior. The threshold is not the same depending on the line, some lines were slower or smaller than others (threshold between 0.6 and 0.3). The validation of these thresholds was performed through ground truthing, contributing to the enhancement of classification, particularly in cases where performance may be suboptimal for certain lines.

The distinction between Static Bends and Head-Cast is determined by applying a threshold to the head velocity. If bending occurs over ‘n’ time steps, the motion velocity normalized by the length of ‘P%’ of those ‘n’ time steps must be below ‘p’ times the mean head velocity of the larva before the stimulation. The values of the two thresholds, ‘P’ and ‘p’, are line-specific, contingent upon the statistical characteristics of the velocity for that particular line; some lines are slower than others.

We computed the cumulative probabilities of actions (Stop, Hunch, Back-up) during five seconds after stimulus onset (as described in Masson *et al,* 2020), only in larvae that were tracked during the entire time window. For actions that occur during baseline locomotion and at high frequency (Crawl and Head Cast), we computed the mean probability over five seconds after stimulus onset and over three seconds starting one second after stimulus onset. We corrected the mean probabilities for these actions by the mean probability computed over twenty seconds of recording prior to stimulus onset. For optogenetic activation of the mechanosensory neurons, because the dynamic of the response was different to that of air-puff experiments, the time window used for computing the cumulative probability of Hunching was two seconds after the stimulus onset and that for bending probability was ten seconds after stimulus onset, by which time bending probability reached baseline levels in larvae fed on standard food. Transition probabilities were computed as the frequency of transition from one action to another over five seconds after stimulus onset, for larvae that were tracked throughout this entire duration.

### Food intake quantification

In order to measure food intake, we quantified the amount of fluorescent food inside the digestive tract of larvae that were allowed to feed ad libitum for 15 min on fluorescent feeding media. To this end, Rhodamine B (Sigma R6626-25G) was diluted in different feeding media (water, yeast extract, sucrose solution or normal food medium, see in media section) to a final concentration of 0.20 µmol/L. 0.8 mI L of each food medium was poured on top of a circular Whatman paper (Fisher scientific, Cytiva 1001-045) placed into a petri dish. 10 larvae were placed into each petri dish. After 15 min of ad libitum feeding on the fluorescent medium, larvae were collected, rinsed in ethanol and in water, and immediately mounted under a coverslip for imaging. Intact larvae were imaged thanks to a fluorescent binocular Zeiss Discovery.V12. The surface of the digestive tract stained by the fluorescent dye was quantified for each larva thanks to a custom-made Fiji script. This script quantified the number of pixels whose intensity was above background.

### Sucrose preference

To measure the preference of larvae between a sucrose-containing agar and a water-containing agar, we performed a place-preference assay. To this end, we prepared petri dishes filled with 0.3% agar diluted in water, divided each agar in two halves, and transferred one half into a new petri dish. Then, we filled the missing half in each petri dish with 0.3% agar diluted in a 20% sucrose solution. Therefore, each petri dish finally contained one side with 20% sucrose agar and one side with agar only.

After cooling down, about 20 third instar larvae were put on the midline of a petri dish and the dish was imaged every 30 s in a behavioral tracker. The number of larvae on each side was then counted over time, excluding the larvae touching the limit between the two media. The preference index was then calculated as: PI = (Ns - Na)/Ntot, where Ns is the number of larvae on the sucrose side, Na the number of larvae on the agar side, and Ntot the total number of larvae.

### Carbohydrate measurements

To measure carbohydrates concentrations inside larval hemolymph, we combined and adapted different methods already published^68,69^. Glucose measurements - Groups of 5 third instar larvae were rinsed in water and placed on a parafilm layer and their cuticle was cut with forceps. 2 µL of the bleeding hemolymph was collected from each group, and 1 µL of 0.05 g/L N-phenylthiourea diluted in PBS was added to avoid darkening of the samples. Samples were heat-inactivated by a 10 min incubation at 90°C and centrifuged 10 min at 10 000 rpm. 1 µL of the supernatant was then mixed with 4 µL of glucose assay kit (Sigma GAHK20-1KT) and incubated for 1 h at 37°C. Absorbance at 340 nm was measured against the blank thanks to a NanoDrop following the manufacturer’s instructions. Hemolymph glucose concentration was finally calculated thanks to a standard curve of glucose concentration.

Trehalose measurements - Groups of 10 third instar larvae were rinsed in water and placed on a parafilm layer and their cuticle was cut with forceps. 3 µL of the bleeding hemolymph was collected from each group, and 97 µL of 0.05 g/L N-phenylthiourea diluted in trehalase buffer (Tris pH 5.5 5 mM, NaCl 137 mM, KCl 2.7 mM) was added to avoid darkening of the samples. Samples were heat-inactivated at 70°C for 10 min, centrifuged for 10 min at 10 000 rpm, and 5 µL of the supernatant were either mixed to 5 µL of trehalase (Sigma T8778-1UN) diluted in trehalase buffer (described above) to a 500 dilution. Samples were incubated at 37°C overnight. 1 µL of each sample was then mixed with 4 µL of glucose assay kit (Sigma GAHK20-1KT) and incubated for 1 h at 37°C. Absorbance at 340 nm was measured against the appropriate blank thanks to a NanoDrop following the manufacturer’s instructions. Trehalose concentration was finally calculated thanks to a standard curve of glucose and trehalose concentrations and by additionally subtracting the concentration of glucose in the sample without trehalase.

### Histochemistry labeling

To determine the neurotransmitter identification in the interneurons, immuno-labeling was performed from the split lines or Gal4 lines crossed to UAS-myr::GFP, or LexA lines crossed to LexAop-myr::GFP. The VNC was dissected out from 3rd instar larvae and fixed with 4% PFA for 45 min at room temperature. After rinsing in PBS, ten minutes permeabilization in PBS-T and two hours blocking in PBS-T-BSA 1%, the CNS preparations were incubated at 4°C (one to three nights) in the first antibodies raised against neurotransmitter and GFP in PBS-T. Then they were incubated at 4°C (one to two nights) in fluorophore-coupled secondary antibodies in PBS-T raised against species of the first antibodies. After rinsing, the preparations were mounted in an anti-bleaching mounting medium (SlowFade Gold, ThermoFisher S36939) under a cover slip. The confocal images were captured with a Leica SP8 confocal laser microscope. Alexa Fluor 488 was excited with a laser light of 488 nm, Cy3 with a laser light of 561 nm, Alexa Fluor 647 with a light of 633 nm wavelength.

### Neuropeptide receptor characterisation

In order to characterize the expression pattern of sNPFR and NPFR in the circuit, expression of two genetically encoded reporter proteins of two different colors was targeted to two different subsets of neurons. To this aim, a T2A Gal-4^48^ or LexA (for sNPFR and NPFR respectively) was used to express LexAop-jRGeco1a or UAS-GCaMP6s (for sNPFR and NPFR respectively) under the control of the promoter of the gene coding for that receptor, thus tagging all neurons which express the receptor transcript. A second genetic driver (LexA for sNPFR and Gal4 for NPFR) was used to individually label target neurons of the circuit with a second reporter protein (UAS-GCaMP6s for sNPFR and LexAop-jRGeco1a for NPFR). The VNC was then dissected and stained with antibodies as described in the previous section.

To the best of our knowledge, no clean or sparse LexA line exists to selectively target the Handle-B and Griddle-2 inhibitory interneuron. For sNPFR expression in Griddle-2, the LexA line L55C05 used targets many neurons in addition to Griddle-2. Griddle-2 could nevertheless be identified by comparing the cell body position and projections in the cross-section of the anterior abdominal segments of the CNS of the R55C05 LexA line and the sparse R55C05 GAL4 line that selectively labels griddle-2 in the VNC (see Extended Data Fig. 7B).

For Handle-B we used a L60E02-LexA line for which a neuron with a cell boy in the midline resembling Handle-B is part of a very dense expression pattern. In order to confirm that the candidate enron was indeed Handle-B, we used the selective split-Gal4 line GMR_SS00888 that specifically labels only Handle-B neurons. We expressed two reporters of different colors (LexAop-GCaMP6s and UAS-Chrimson-mCherry under the control of GMR_SS00888 and L60E02 in the same larva. The colocalization confirmed the line 60E02-LexA to target Handle-B (Extended Data Fig. 7C).

### In vivo imaging of intact larvae

Because opening the cuticle might affect the larval internal state, we developed a simple preparation for the imaging of intact larvae. For this purpose, third instar larvae were rinsed in water, and mounted between a 2 cm circular coverslip and a custom-made device that delivers mechanical stimulations in low melting point agarose 4% (melted in phosphate buffer saline), ventral side facing up. Larvae were gently squeezed in this position until agar cooled down, so that the ventral nerve cord could be imaged through the cuticle.

All Gal4 and LexA drivers used for *in vivo* imaging are listed in the figure legends. The imaging plane was restricted to the location of the projections of neurons of interest, in particular when sparse lines that lacked specificity towards a unique neuronal type were used (R22E09).

Mechanical stimulations were generated by a waveform generator (Siglent sdg1032x) connected to a quick-mount extension actuator (Piezo Systems, Inc.), which was embedded in the sylgard-coated recording chamber (Sylgard Silicone Elastomer, WPI). The stimulation was set at 1000 Hz, with an intensity of 1 to 20 V applied to the actuator. The amplitude of the acceleration produced by the actuator was measured thanks to a triple axis accelerometer (Sparkfun electronics ADXL313) connected to a RedBoard (Sparkfun electronics) and bound to the sylgard surface thanks to high vacuum grease. Acceleration was 1.14 m.s^-2^ at 20 V, and 0.61 m.s^-2^ at 10 V. Mechanical stimulations were precisely triggered by the Leica SP8 software thanks to the Leica “Live Data Mode” and to a trigger box branched to the scanning head of the microscope. A typical stimulation experiment consisted in 5 s of recording without stimulation, then 5 s of stimulation, and 5 s of recording in the absence of stimulation.

For optogenetic activation during in vivo imaging, larvae were mounted in the dark, with the least intensity of light possible in the room, to avoid nonspecific activation of the targeted neurons. Optogenetic stimulation of CsChrimson was achieved by a 617-nm wavelength LED (Thorlabs, M617F2), controlled by a LED driver (Thorlabs, LEDD1B) connected to the waveform generator, and conveyed through a Ø 400 µm Core Patch Cable (Thorlabs) to the imaging field. Optogenetics stimulations were triggered at 50 Hz, 50% duty during 1 s, concomitantly to mechanical stimulations thanks to the waveform generator. Irradiance was measured at the level of the imaging field at 500 µW using a PM16-130 THORLABS photometer.

Imaging was achieved with 1-photon or 2-photon scanning Leica SP8 microscope, at 200 Hz, with a resolution of 512 x 256 pixels or 512 x 190 pixels. The rate of acquisition was 1 frame/s or 2 frames/s depending on the experiment. For Basin-2 recordings, the stimulation was repeated 5 times with resting intervals of 60 s in order to calculate a frequency of response. Recordings where the dF/F averaged over the whole stimulus duration did not exceed 10% were considered as failed responses. Optogenetic experiments were conducted with 2-photon imaging.

When the projections of the neurons were not visible before stimulation (imaging of Handle-b upon NPFR knockdown), we used resonance scanning with 10 line accumulation and 6 frame averaging to increase the signal. This resulted in one image being taken each second and in the dF/F being lower than usual. With these settings one recording was acquired per larva.

Neuronal processes were imaged in the VNC at the axonal level and fluorescence intensity was measured by manually drawing a region of interest (ROI) in the relevant areas using custom Fiji macros. Data were further analyzed using customized MATLAB scripts. F0 was defined as the mean fluorescence in the ROI during baseline recording, in the absence of mechanical stimulus or optogenetic activation. ΔF/F0 was defined at each time step t in the ROI as: ΔF/F0 = (F(t) - F0)/F0.

For recording the baseline activity level of NPF neurons, one frame was recorded each second for 20 seconds and, for each larva, the frame showing the most intense fluorescence in the neuronal projections was used to evaluate its raw fluorescence level.

For imaging neurons in different feeding conditions, the effect of food treatment on the expression level of fluorescent reporter proteins was assessed by expressing GFP in the chordotonal neurons. Fluorescence was measured after exposing larvae to different food treatments.

## Statistical analysis

### Locomotion analysis

For all boxplots and histograms pairwise Mann-Whitney tests were used to evaluate significant differences between larva groups, with Bonferroni correction for multiple comparisons. Significance was illustrated according to the p-value by asterisks in histograms (*:<0.05, **:<0.01, ***:<0.001, ****:<0.0001) and by pairs of colored semicircles in boxplots, the left always corresponds to the group with highest mean value. Non-significant tests were omitted for visual clarity.

### Behavior probabilities

Chi² tests were used for statistical analysis of behavioral and transition probabilities.

To assess the effects of different states/neuronal manipulations on Head Casting in response to air-puff, we calculated an estimator designed to identify the emergence of behaviors at the population scale. We aim to determine the probability induced by the stimulation, so we need to subtract the stationary probability without the stimulus. We calculate the probability of larvae bending 5 seconds after the stimulus, on tracking larvae throughout this time window (denoted as *p*_*A*_). For the probability of bending before the stimulus, we consider all larvae tracked continuously for 20 seconds prior to the stimulus, between 30 and 50 seconds (denoted as *p*_*B*_). A probability is defined as *p*_*k*_ = *N*_*k*_/*N*_*k,all*_ with *k* ∈ {*A*, *B*} and *N_k,all_* the total number of larva taking to compute probabilities. In order to quantify the effect of the stimulus we defined Χ = *p*_*A*_ − *p*_*B*_ as the difference in the ratio after and before the stimulus.

In order to compare test line our estimator was defined as

Θ(*p*, *q*) = χ(*p*) − χ(*q*) with *p* and *q* the ratios of the lines and the control respectively. Θ(*p*, *q*) takes value in [−1,1]. The null hypothesis is Θ(*p*, *q*) = 0, if there are no differences between the line tested and the control. Positive or negative values indicate an effect of neuron silencing when compared to the control.

We use numerical simulations to conduct a statistical test where {*p*_*k*_, *q*_*k*_} are generated from a hypergeometric distribution, *X* ∼ *Hypergeometric*(*N*_*k,all*_, *N*_*k*_, 1). We perform *N_all_* = 10^3^ repetitions, computing Θ(*p*, *q*) each time. The p-value is determined by the number of instances when the hypothesis is not verified, divided by the total number of repetitions, (pseudo-code in Jovanic *et al*, 2016^35^).

Note that this estimator, Θ(*p*, *q*), has the advantage of being able to detect the non-synchronous emergence of a behavior at a population scale. For example, head casting can either emerge as an immediate response to the puff or as the second response after Hunching. The statistics of start time of Head Casting is thus widely distributed at the population scale. Time evolution of the instantaneous ratio of larva performing bending would not exhibit a strong increase after stimuli because larvae are not all going to bend immediately after stimuli. Θ(*p*, *q*) by accumulating events during a time window allows efficient detection of a behavior even if it is widely distributed in time.

### Calcium imaging

All data in line plots are presented as mean ± SEM. Violin plots show the first and third quartiles, the average of all recordings as a white line and the median as a white dot. Comparisons of the data series between two conditions were achieved by a two-tailed unpaired t-test. Comparisons between more than two distinct groups were made using a one-way ANOVA test, followed by Bonferroni pairwise comparisons between the experimental groups and their controls.

### Mathematical model

We reproduced and extended the rate-based system model of the circuit that was published in a previous publication^35^. The circuit is described as a rate model with a connection matrix derived from the larva connectome. Each neuron population (mechano-ch, iLNa, iLNb, fbLN-Ha and fLN-Hb) was modeled by a single node (Fig. 3C-E). The dynamics read:

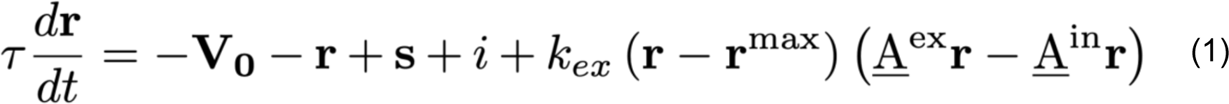

with **τ** representing the vector of the characteristic time constants of the neurons, **r** the rate vector, **V_0_** the threshold vector, **s** the sensory stimulus input vector, **i** the vector of inputs from other brain regions, k_ex_ a sensitivity factor to overall input, **r**^max^ the maximal rate vector, A^ex^ and A^in^ respectively the excitatory and inhibitory coupling matrices.

Values in <u>A</u>^in^ and <u>A</u>^ex^ were directly extracted from synaptic counts (see Jovanic *et al.,* 2016).

In order to represent the variety of stimuli larvae are subjected to, and thus the variety of behavior they elicit, we follow the approach in Jovanic *et al.*, 2016 and vary the connection strength between Mch and iLNa populations. In the original paper, the connection strength between Mch and iLNb populations is also varied. We decided to fix the value of this connection strength at 2 in order to reduce the number of parameters to explore.

We used the *solve_ivp* routine of the integration package from SciPy, which internally calls the LSODA solver, able to switch between the Adams method and the BDF method, based on the stiffness of the equation. We used relative and absolute tolerances of 10^-3^. Additionally, the solution vector is constrained to stay positive. This is obtained by replacing r by max(r, 0), and the components of r’ by those of max(r’, 0) whenever the corresponding component of r is smaller than 10^-9^.

The behavior is defined based on the end state of the network and, more specifically, the rates of the neurons B1 and B2. We used a k-means clustering with k=2 to separate the values of the output neurons for an ensemble of simulations corresponding to connection strengths spanning [0.5, 1.5] between MCh and iLNa, and [1.5, 2.5] between iLNa and iLNb. The output activations cluster strongly in a coactive state (rate(B1) > 0, rate(B2) > 0) corresponding to bends, and a monoactive state (rate(B1) > 0, rate(B2) = 0) corresponding to Hunches. The two behaviors can also be distinguished by the single scalar rate(B2)/rate(B1), which is positive for bends and zero for Hunches. This ratio is sometimes plotted instead of the discrete category.

We explore two models for neuromodulation, which modify the behavior output of the network without altering its connectivity.

The first model hypothesizes that in the sucrose state, Hb receives an additional input current, modeled by a nonzero entry to the vector **i**. We show that as this input current increases, the range of stimuli evoking bends increases, allowing us to claim that an additional input current to Hb increases the likelihood of bends. We further fix the value of the input current to 10 a.u. in the sucrose state and 0 a.u. in the fed state, to perform silencing analyses.

The second model hypothesizes that in the sucrose state, the maximum rate for the iLNa neuron population is decreased, representing a saturation of the response to external stimuli. We show that as the r_max_ parameter for iLNa decreases, the range of stimuli evoking bends increases, consistent with experimental observations and once again despite the use of arbitrary units. For silencing analyses, we define the sucrose state as r_max_ = 18 a.u. and the fed state as r_max_ = 20 a.u. Finally, we also consider a model combining both hypotheses. In this model, every combination of a decrease in r_max_ for iLNa and an increase in input current to Hb results in more bend. For silencing analyses, we fix the exact values of those parameters. In the combined model, we define the sucrose set by setting each parameter to the value defining the sucrose state in the single hypothesis models.

We provide here the list of parameters used in the simulations.

|  |  |
| --- | --- |
| $k_{\text{ex}}$ | 2.5 |
| $i$ | [0, 0, 0, 0, 0, 0, 0] in fed state<br>[0, 10, 0, 0, 0, 0, 0] in sucrose state for the combined model |
| $r_{\max}$ | [20, 20, 20, 20, 20, 20, 20] in fed state<br>[20, 20, 20, 20, 20, 20, 18] in sucrose state for the combined model |
| $V_0$ | [0, 20, 20, 20, 20, 20, 20] |
| □ | [1, 35, 35, 35, 35, 35, 35] |

The excitatory and inhibitory matrices read

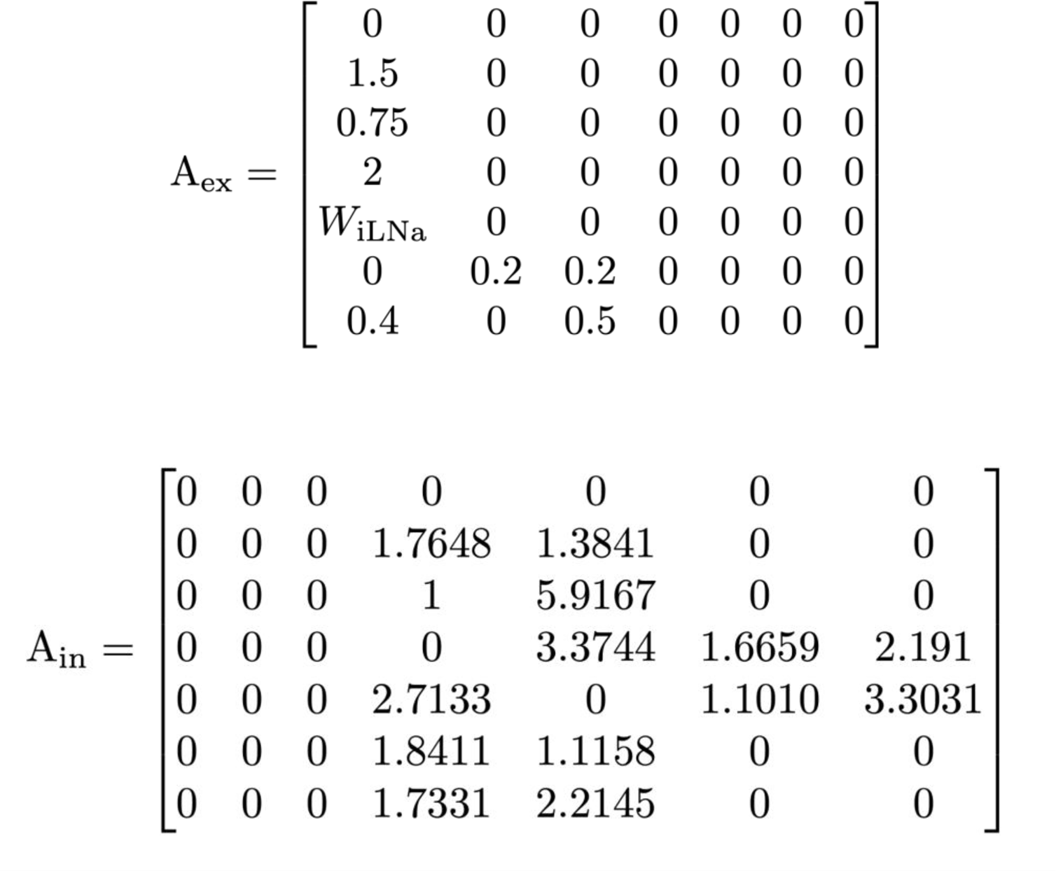

## Extended Data Figures and Legends

**Extended Data Fig. 1.**
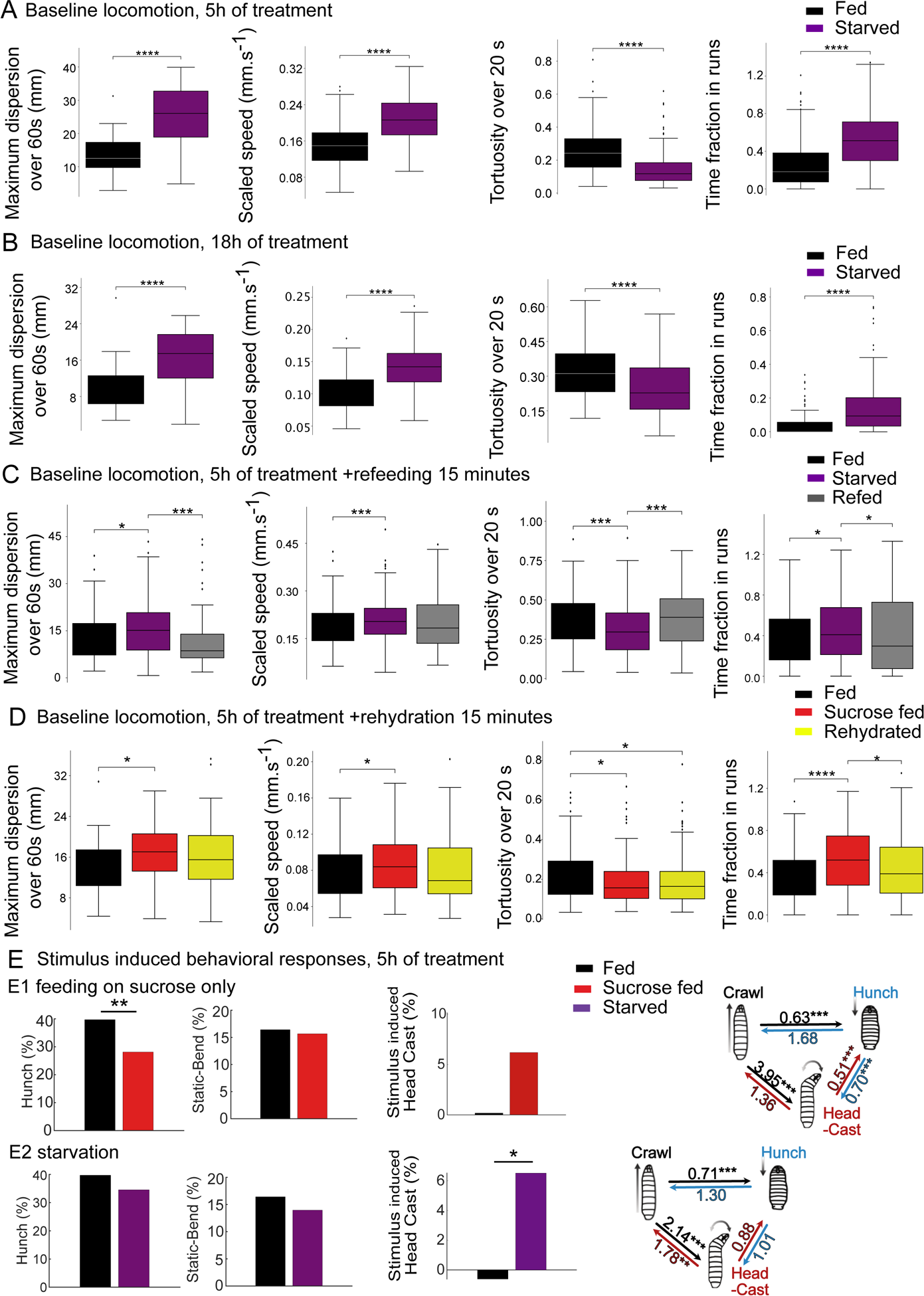
Locomotion of food-deprived larvae. A-D: Analysis of larval locomotion in different feeding states. A-B: Larvae starved for 5h (A, n = 283, 211, 309 larvae) or 18h (B, n = 173, 141, 139 larvae) disperse further, at a higher movement speed, allocating more time to crawling (time fraction in runs) with lower-tortuosity trajectories compared to larvae fed on a standard diet. C: Refeeding larvae on standard food for 15 min after a period of 5h starvation rescues the normal locomotion as dispersion distance, time allocation to crawling and crawling speed are no longer significantly increased, nor the trajectories’ tortuosity decreased compared to larvae fed on a standard diet (n= 244, 160, 223 larvae) D: Rehydration by putting larvae for 15 minutes on water after 90 minutes of feeding of sucrose restores normal locomotion similar to larvae fed on standard food, since dispersion distance, time allocation to crawling and movement speed are no longer significantly increased, nor the trajectories’ tortuosity decreased. (Mann-Whitney test, ****: p<0.0001, ***: p < 0.001, **: p < 0.01, *: p < 0.05) (n = 102, 109,123 larvae) E-F: Behavior in response to air-puff during the first five seconds upon stimulus onset, Behavioral and transition probabilities upon 5h of sucrose feeding (E) and starvation (F) (n=233-301 larvae, ***: p < 0.001, **: p < 0.01, *: p < 0.05).

**Extended Data Fig. 2.**
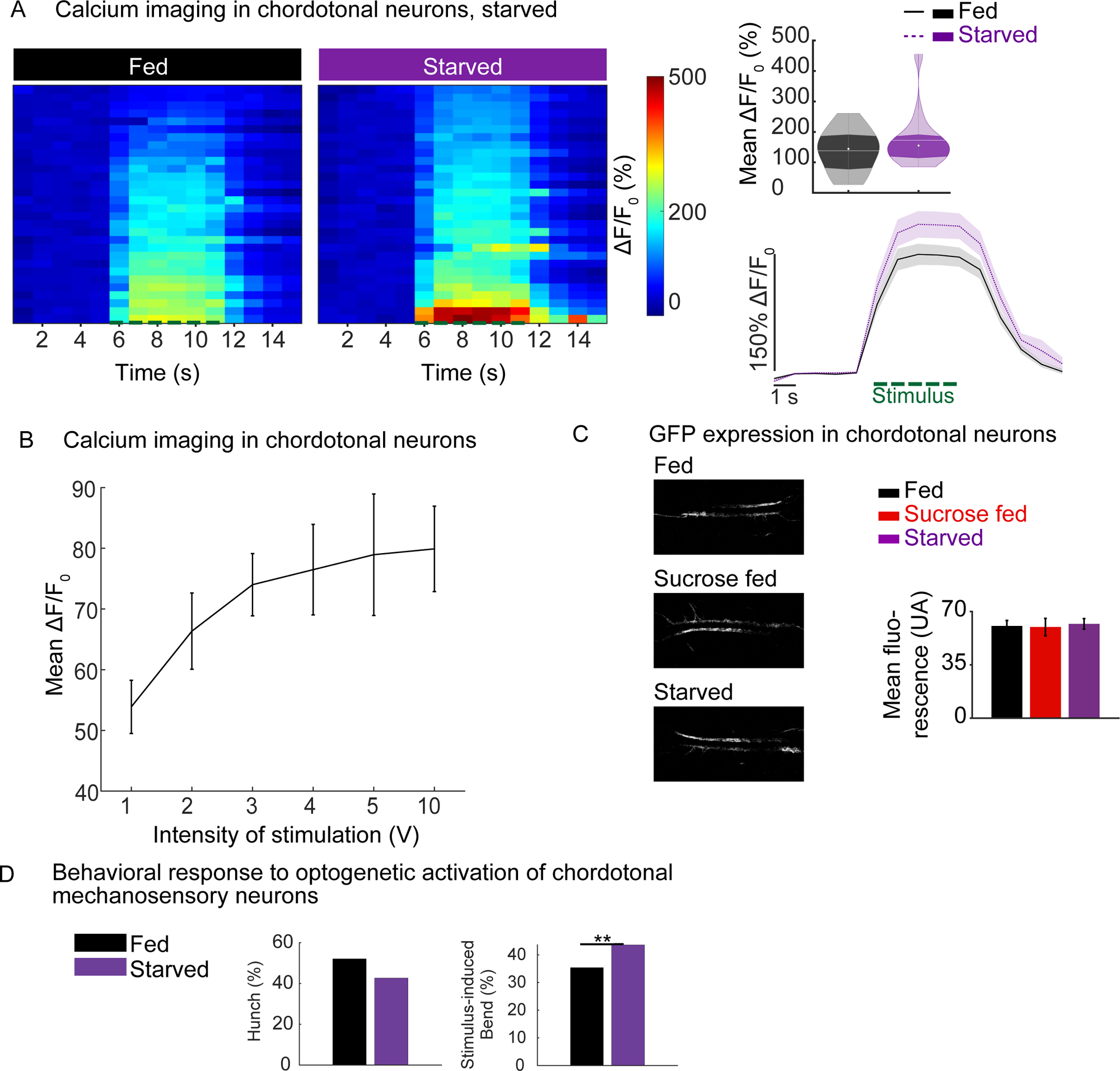
Feeding state dependent bias in sensorimotor responses does not come from the modulation of sensory neurons. A: Calcium responses to mechanical stimulations at 5V in chordotonal neurons in fed and starved larvae (R61D08-Gal4/UAS-GCaMP6s). Left panel: calcium responses of chordotonal neurons from different individuals fed on different diets. Lower right panel: mean calcium response of chordotonal neurons over time. The green dashed line corresponds to stimulus duration. Upper right panel: calcium response averaged during the stimulus. White line represents the mean, white dot represents the median. Stimulus-induced activity of chordotonal mechanosensory neurons is not significantly increased in starved animals as compared to larvae fed on a standard food (n = 10 larvae, 1 trial per larva, t-test p = 0.092). B: Calcium responses of chordotonal neurons to different intensities of mechanical stimulations, in larvae fed on standard food medium (n = 5 larvae). C: GFP expression levels in chordotonal neurons R61D08>GFP larvae fed on different food media: standard food, 20% sucrose and water (n = 4 larvae per condition). D: Optogenetic activation of cho in starved larvae. Hunch is cumulative probability during the first 2 seconds from stimulus onset. Bend is the mean probability during the first 10 seconds from stimulus onset, corrected by 40 seconds of recording prior to the stimulus. Hunching and stimulus-induced Bending (n = 192 larvae for fed, 157 for starved, p = 0.080 for hunch, 0.003 for bend).

**Extended Data Fig. 3.**
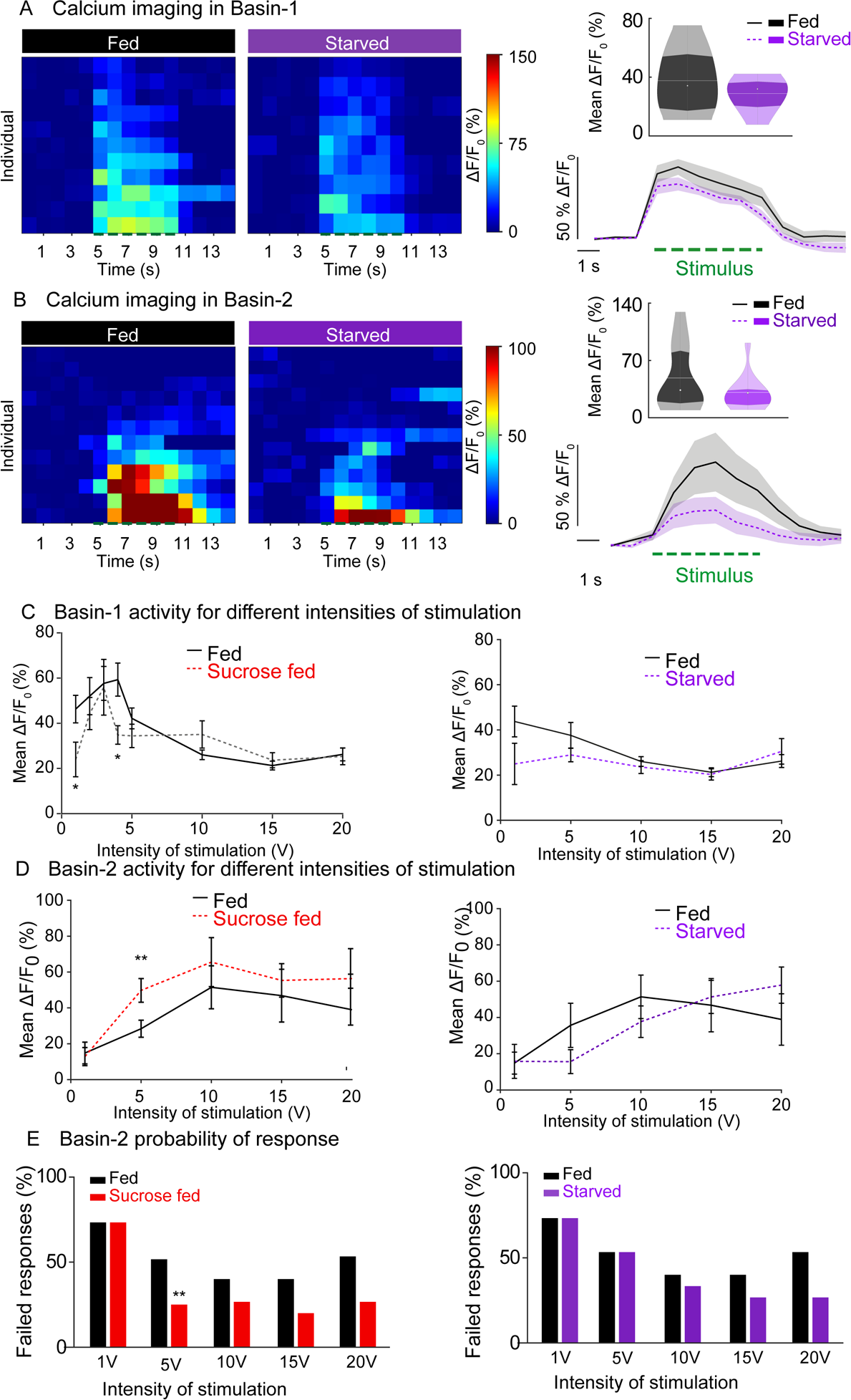
Effect of starvation and different stimulus intensity on responses of projection neurons to mechanical stimulations. A: Basin-1 calcium responses to mechanical stimulations in fed and starved larvae (20B01-lexA; LexAop-GCaMP6s, UAS-CsChrimson-mCherry). Left panel: calcium responses of Basin-1 from different individuals fed on different diets. Lower right panel: mean calcium response of Basin-1 over time. The green dashed line corresponds to stimulus onset. Upper right panel: mean calcium response averaged during the stimulus. White line represents the mean, white dot represents the median. Stimulus-induced activity of Basin-1 neurons is similar in starved animals compared to larvae fed on a standard food (n = 11/9 larvae, 1 trial per larva, t-test p = 0.2892). B: Basin-2 calcium responses to mechanical stimulations in different states (SS00739/UAS-GCaMP6s). Left panel: calcium responses of Basin-2 from fed and starved animals. Lower right panel: mean calcium response of Basin-2 over time. The green dashed line corresponds to stimulus onset. Upper right panel: mean calcium response averaged during the stimulus. White line represents the mean, white dot represents the median. Stimulus-induced activity of Basin-2 is not significantly decreased in starved animals compared to larvae fed on a standard food (n = 12/13 larvae, 1 trial per larva, t-test p = 0.1804). C: Basin-1 calcium responses for different intensities of stimulation, in larvae fed on standard food, on sucrose only or completely starved larvae (n = 8-20 larvae, 1 trial per larva). Average neuronal activities over time for different intensities of stimulation are plotted. D: Basin-2 calcium responses for different intensities of stimulation, in larvae fed on standard food, on sucrose only or completely starved larvae(n = 11-13 larvae, 1 trial per larva, t-test **: p < 0.01). Averaged neuronal activity over time for different intensities of stimulation are plotted. E. percentage of failed responses comparison Chi-2 test p = 0.0027).

**Extended Data Fig. 4.**
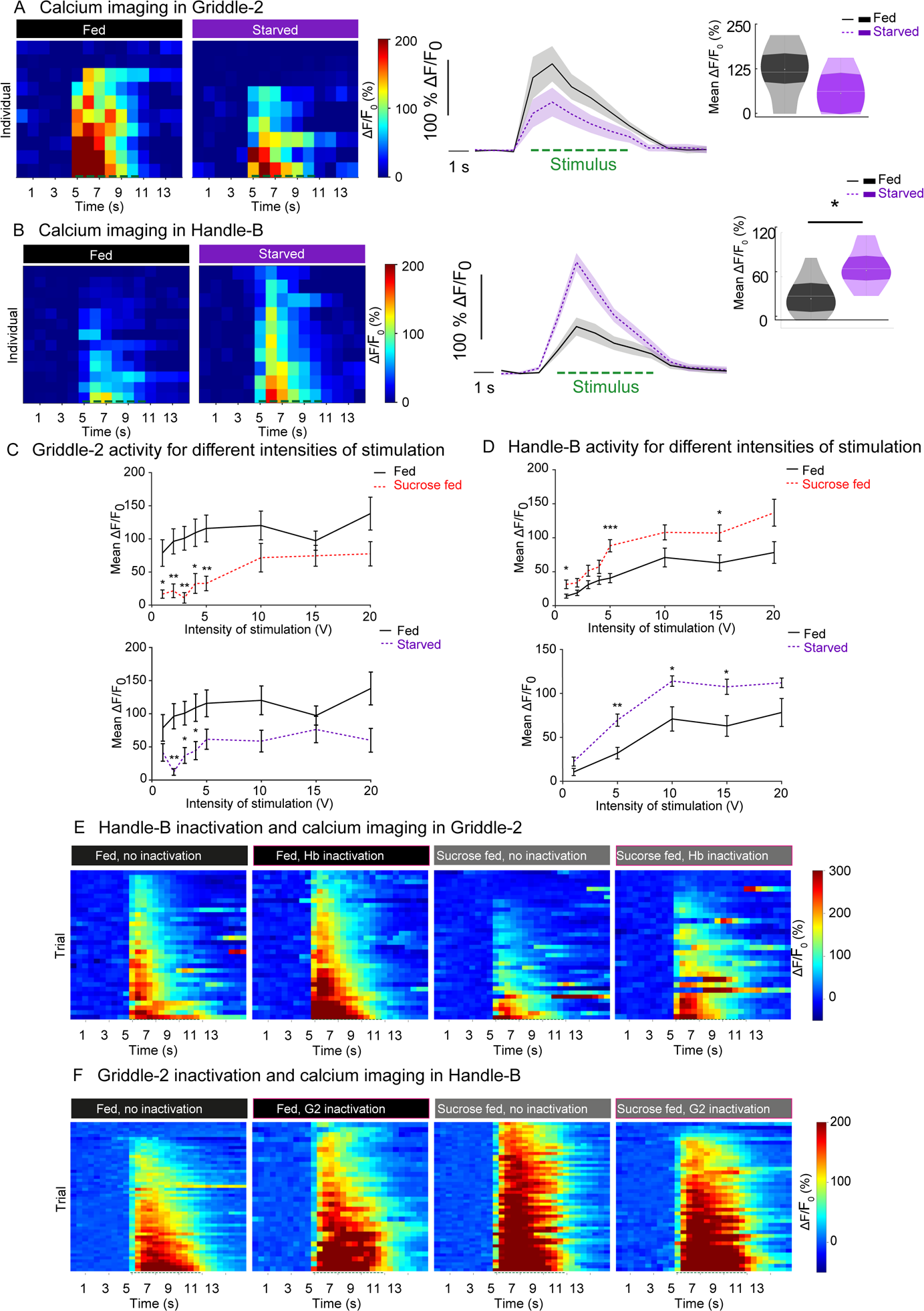
Two interneuron subtypes are oppositely modulated by the feeding state. A: Griddle-2 calcium responses to mechanical stimulations in fed and starved larvae (SS00918/UAS-GCaMP6s). Left panel: calcium responses of Griddle-2 from different individuals fed on different diets. Lower right panel: mean calcium response of Griddle-2 over time. The green dashed line corresponds to the stimulus Upper right panel: mean calcium response averaged during the stimulus. White line represents the mean, white dot represents the median. Stimulus-induced activity of Griddle-2 neurons is not significantly decreased in starved animals compared to larvae fed on a standard food (n = 10/9 larvae, 1 trial per larva, t-test p = 0.0979). B: Handle-b calcium responses to mechanical stimulations in different states (SS00888/UAS-GCaMP6s). Left panel: calcium responses of Handle-b from fed and starved animals. Lower right panel: mean calcium response of Handle-b over time. The green dashed line corresponds to stimulus onset. Upper right panel: mean calcium response averaged during the stimulus. White line represents the mean, white dot represents the median. Stimulus-induced activity of Handle-b is significantly increased in starved animals compared to larvae fed on a standard food (n = 10/13 larvae, 1 trial per larva, t-test p = 0.019). C: Griddle-2 calcium responses for different intensities of stimulation, in larvae fed on standard food, on sucrose only and starved larvae (n = 9-10 larvae, 1 trial per larva, t-test **: p < 0.01, *: p < 0.05). Average neuronal activities over time for different intensities of stimulation are plotted D: Handle-b calcium responses for different intensities of stimulation, in larvae fed on standard food or on sucrose only.and starved larvae (n = 9-25 larvae, 1 trial per larva, t-test ***: p < 0.001, **: p < 0.01, *: p < 0.05). Responses averaged during the stimulus for different intensities of stimulation are plotted E. Calcium responses in Griddle-2 with (55C05-LexA>LexAop-GCaMP6s SS00888-Gal4>UAS-TNT) or without Handle-b inactivation (55C05-LexA>LexAop-GCaMP6s +/UAS-TNT), in each trial of mechanosensory stimulation (n = 7-9 larvae, 4 trials per larva). F: Calcium responses in Handle with (55C05-LexA>LexAop-TNT 22E09-Gal4>UAS-GCaMP6s) or without Handle-b inactivation (+/LexAop-TNT 22E09-Gal4>UAS-GCaMP6s), in each trial of mechanosensory stimulation (n = 6-10 larvae, 5 trials per larva).

**Extended Data Fig. 5.**
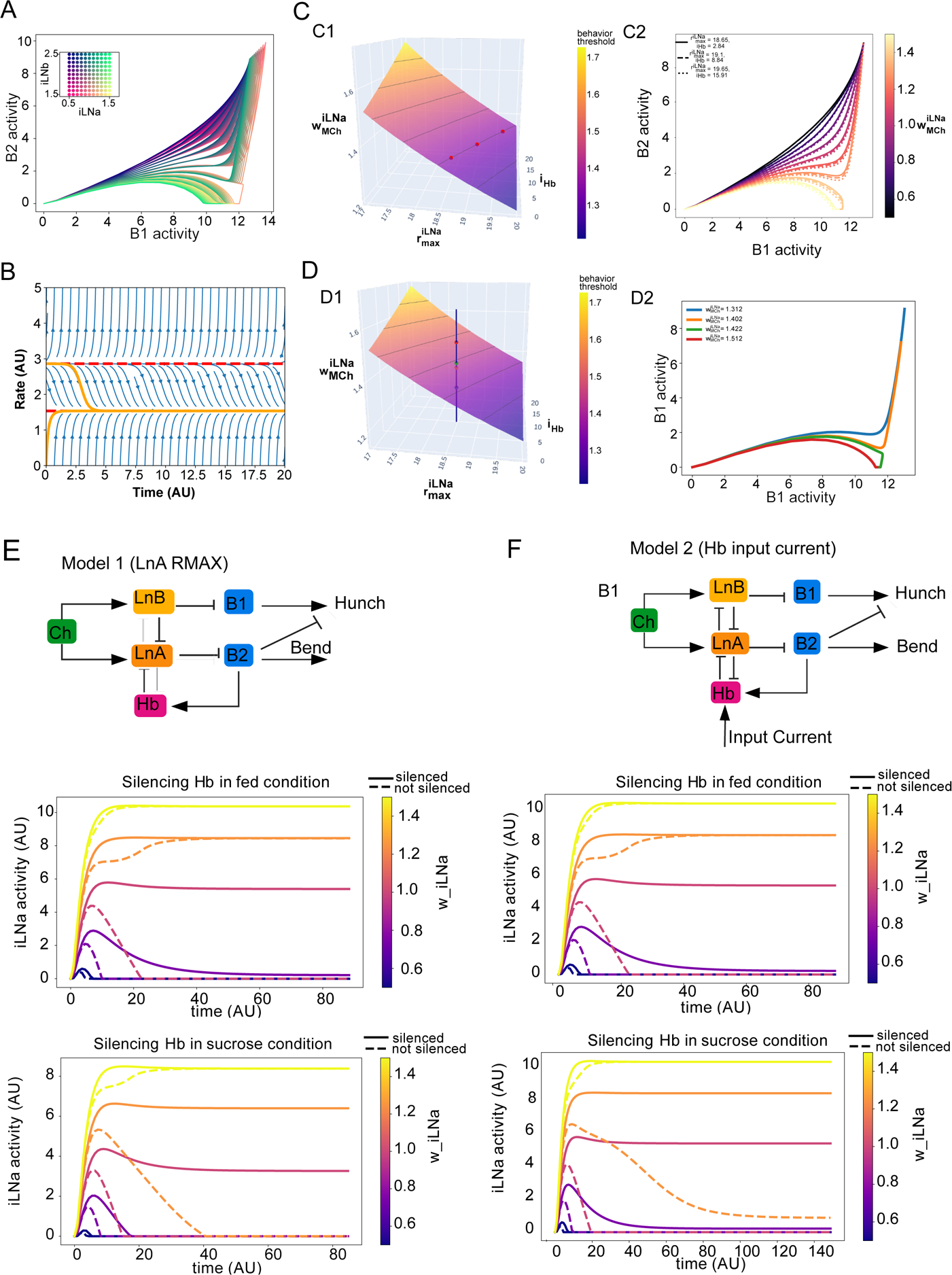
Circuit model. A: State space trajectories of B1 and B2 Dynamics as a function of inputs to LNa and LNb neurons in a simple rate model (as in Jovanic *et al.,* 2016). B: Dynamics of a single model neuron with self inhibition shows two equilibria, only one of which is stable C: Level-set for threshold in combined model. C1. Points on lines at the intersection of the surface and horizontal planes represent sets of parameters with identical behavior response. C2. State-space trajectories of the models parameterized by the three points in C1 do not differ qualitatively D: Combined model D1. Points on a vertical line correspond to the same model parameter, with different inputs to iLNa. Points above the surface converge to the monoactive state, while points below the surface converge to the coactive state. D2.Trajectories corresponding to the points in D1, color-coded. E: Silencing Handle-B in model 1, where the sucrose state is modeled as a decrease in LNa max. F: Silencing Handle-b in model 2 where the sucrose state is modeled as increase input in Handle-b.

**Extended Data Fig. 6.**
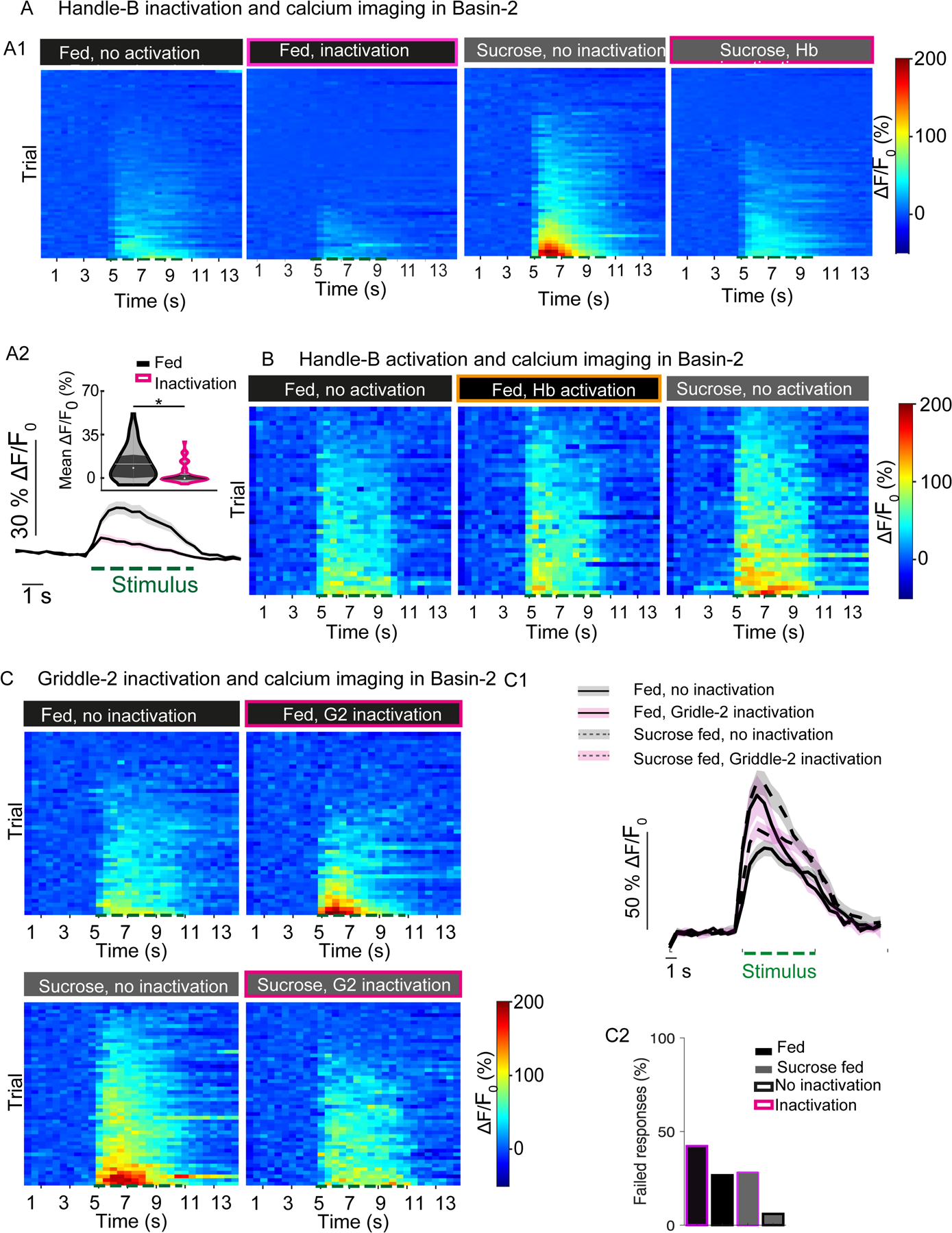
Inhibitory interneurons are required for the feeding state dependent modulation of projection neurons. A1: Calcium responses in Basin-2 with (38H09-LexA>LexAop-GCaMP6s 22E09-Gal4>UAS-TNT) or without (38H09-LexA>LexAop-GCaMP6s +/>UAS-TNT) Handle-b inactivation in each trial of mechanosensory stimulation, in larvae fed on different diets (n = 12-14 larvae, 5 trials per larva). A2: calcium response of Basin-2 averaged during the stimulus. White line represents the mean, white dot represents the median with (38H09-LexA>LexAop-GCaMP6s 22E09-Gal4>UAS-TNT) or without (38H09-LexA>LexAop-GCaMP6s +/>UAS-TNT) (n = 12-14 larvae, 5 trials per larva; ANOVA with post-hoc tests: *: p < 0.05. B: Individual calcium responses in Basin-2 from individuals fed on different diets, with or without optogenetic activation of Handle-b (22E09-Gal4>UAS-CsChrimson::tdTomato 38H09-LexA>LexAop-GCaMP6s) during the first second of mechanical stimulus (n = 8 larvae, 5 trials per larva). In all plots, the green dashed line corresponds to stimulus onset. C: Calcium responses in Basin-2 with (38H09-LexA>LexAop-GCaMP6s 55C05-Gal4>UAS-TNT) or without (38H09-LexA>LexAop-GCaMP6s +/>UAS-TNT) Griddle-2 inhibition, in each trial of mechanosensory stimulation, in larvae fed on different diets (n = 6-10 larvae, 5 trials per larva).

**Extended Data Fig. 7.**
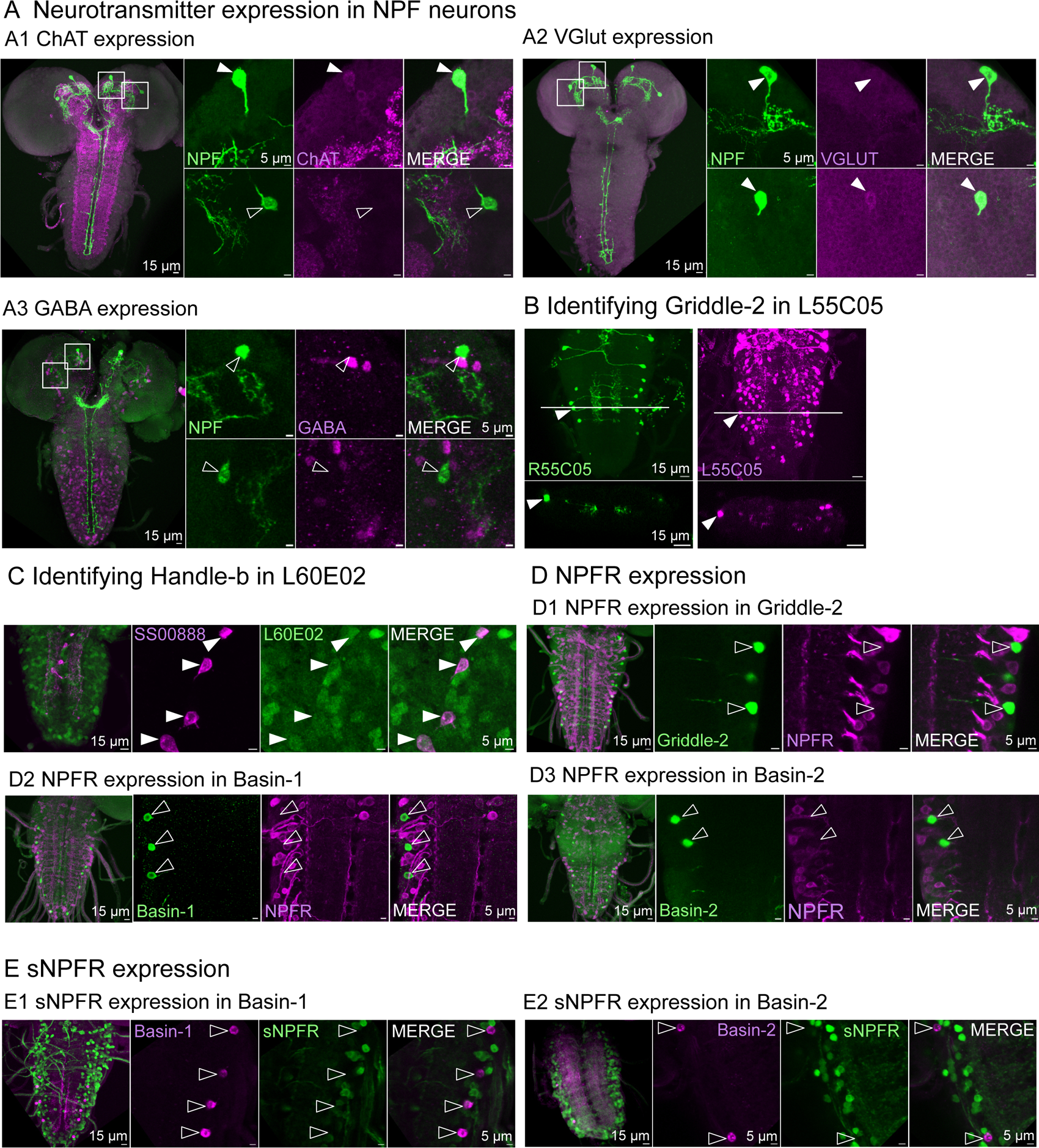
Neurotransmitter identity of NPF neuron and NPF and sNPF receptor expression in neurons of the circuit. A.: Immunohistochemical labeling of NPF neuron neurotransmitters (NPF-Gal4>UAS-GFP) labels both pairs of NPF expressing neurons: dorsomedial (DM) and dorsolateral (DL). Brains were then stained with antibodies against either ChAT as a proxy of acetylcholine (A1), V-Glut as a proxy of glutamate (A2) and GABA (A3). Co-localization shows expression of acetylcholine in the DL NPF neurons and glutamate in both pairs of NPF descending neurons. Both pairs were negative for GABA antibody labeling. B: identifying griddle-2 in 55C05. The position of cell bodies and projections was compared between neurons in R55C05 (GAL4) (previously identified as Griddle-2) and L55C05 (LexA) that had a much broader expression pattern. C: Identification of Handle-b in line L60E02. L60E02 is driving the expression of LexAop-GCamP6s and GMR_SS00888 the expression of UAS-Chrimson-mCherry (60E02-LexA; UAS-Chrimson-mCherry, LexAop GCamP6s>GMR_SS00888). Specific antibodies against GFP and mCherry were used to increase detection sensitivity. Co-localization of jRGeco1a and GCamP6s show that the neuron with the cell body in the midline labeled by the 60E02-LexA is Handle-b. D1. Immunohistochemical labeling for NPFR in Griddle-2. UAS-GCamP6s is expressed in Griddle-2 using the SS_TJ001 split-Gal4 line (green) and LexAop-jRGeco1a is expressed under the control of the NPFR promoter using a T2A-LexA construct (magenta). Antibodies against GFP and dsRed were used to increase detection sensitivity. No expression of NPFR could be detected in Griddle-2. D2-D3: Basin-1 and -2 immunostaining for NPFR expression. UAS-GCamP6s is expressed in Basin-1 or Basin-2 using the R20B01 or GMR_SS00739 driver lines (green) respectively and LexAop-jRGeco1a is expressed instead under the control of the NPFR transcript using a T2A-LexA construct (magenta). Antibodies against GFP and dsRed were used to increase detection sensitivity. No expression of NPFR could be detected in Basin-1 or -2. E: Basin-1 and -2 immunostaining for sNPFR expression. LexAop-jRGeco1a is expressed in Basin-1 or Basin-2 respectively using the L20B01 or L38H09 lines (magenta) and UAS-GCamP6s is expressed under the control of the sNPFR promoter using a T2A-Gal4 construct (green). Specific antibodies against GFP and dsRed were used to increase detection sensitivity. No expression of NPFR could be detected in Basin-1 or -2.

**Extended Data Fig. 8.**
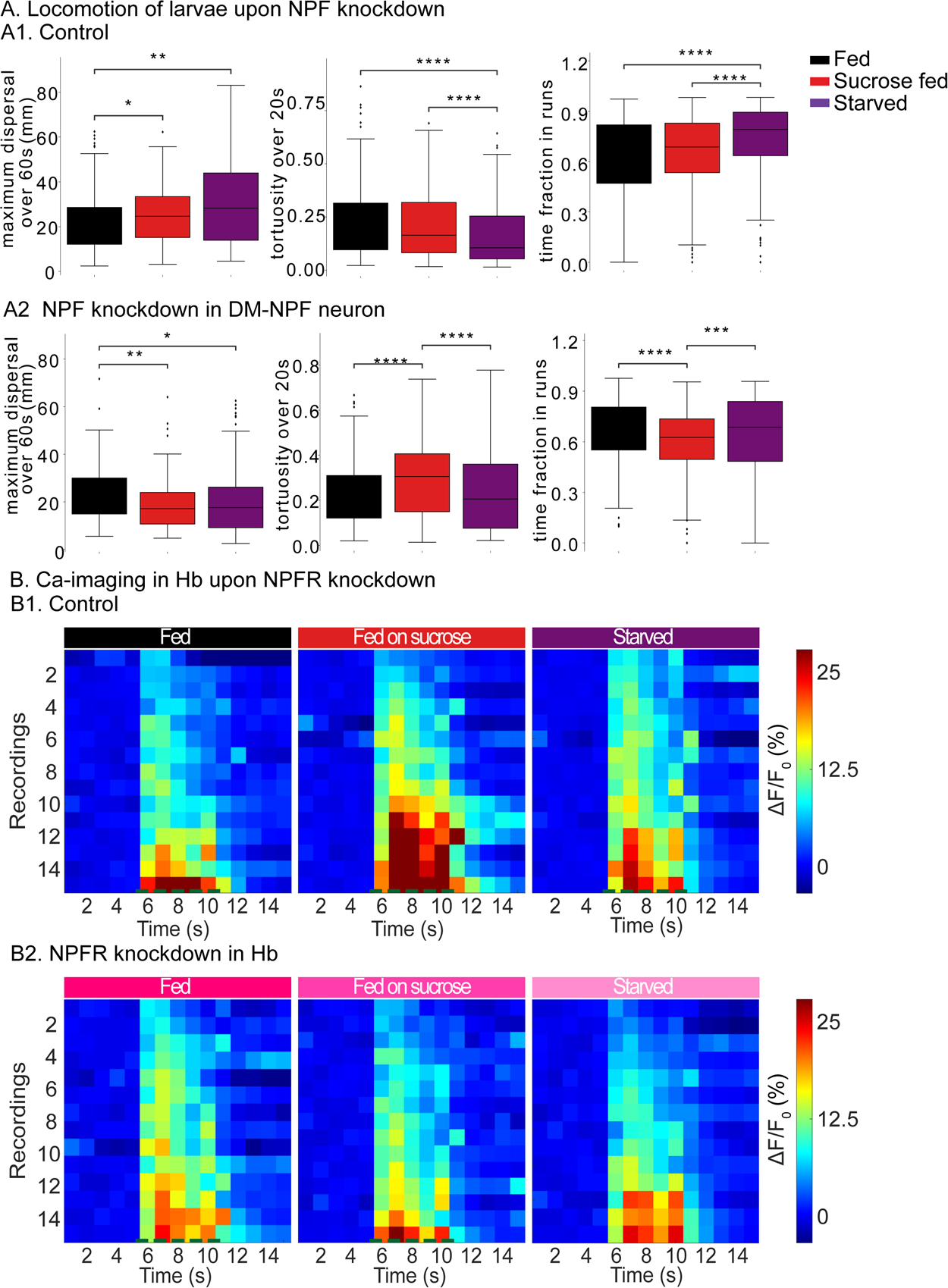
NPF regulates feeding state dependent changes in locomotion and Handle-b activity. A: Analysis of larval locomotion in the absence of sensory stimulus. Top row shows control larvae (A1). Bottom row shows larvae with NPF knockdown in DM-NPF descending neurons (A2). NPF knockdown in DM-NPF descending neuron abolishes differences in locomotion mean between larvae in different states; it abolishes the difference in trajectory tortuosity, time allocation in exploration (time fraction in runs), and dispersal between larvae fed on standard food and starved larvae. In larvae fed on sucrose the increase in exploration compared to larvae fed on standard food was also abolished (Mann-Whitney test, ****: p<0.0001, ***: p < 0.001, **: p < 0.01, *: p < 0.05) B: Calcium responses in Handle-b upon NPFR knockdown (GMR_SS00888>UAS-NPFR-RNAi; UAS-GCamP6s) compared to a control (GMR_SS00888>UAS-GCamP6s). Top row: calcium responses of Handle-b in control larvae fed, fed on sucrose or starved, in each trial of mechanosensory stimulation (n = 15 larvae per condition). Bottom row: calcium responses of Handle-b upon NPFR knockdown in larvae fed, fed on sucrose or starved, in each trial of mechanosensory stimulation (n = 14-15 larvae per condition).

**Extended Data Fig. 9.**
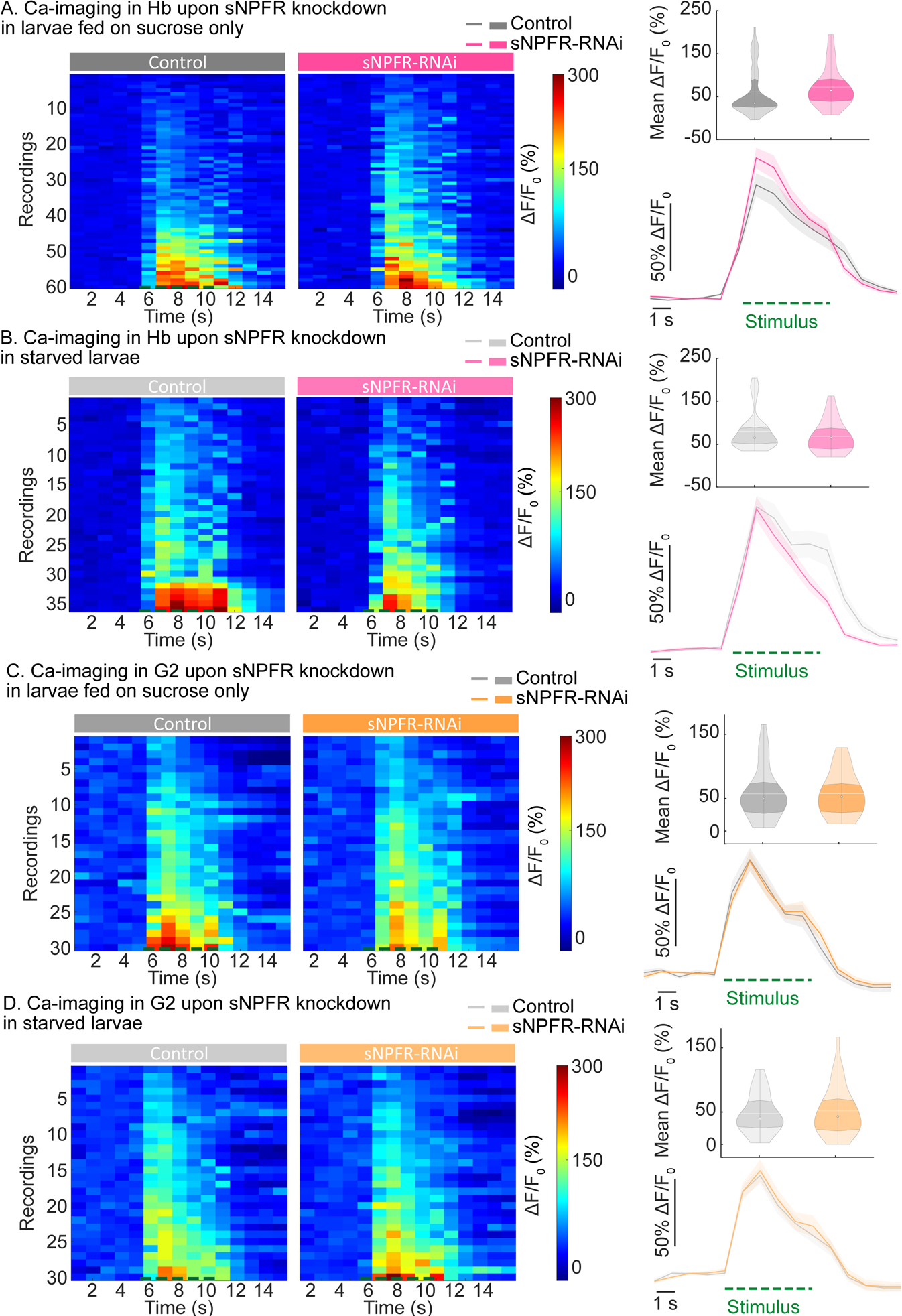
sNPFR knockdown does not impact interneurons responses in larvae fed on sucrose or starved larvae. A: Calcium responses in Handle-b sNPFR knockdown in Handle-b (GMR_SS00888>UAS-GCamP6s; UAS-sNPFR-RNAi) compared to the control (GMR_SS00888>UAS-GCamP6s) in larvae fed on sucrose only (top row) or starved (bottom row). Left panel: calcium responses of Griddle-2, in each trial of mechanosensory stimulation. Lower right panel: mean calcium response of Handle-b over time. The green dashed line corresponds to stimulus onset. Top right panel: response averaged during the stimulus. White line represents the mean, white dot represents the median. sNPFR knockdown does not influence the stimulus-induced activity of Handle-b in larvae fed on sucrose only or starved (n = 9-15 larvae, 4 trials per larva). B: Calcium responses in Griddle-2 with (SS_TJ001>UAS-GCamP6s; UAS-sNPFR-RNAi) or without (SS_TJ001>UAS-GCamP6s) sNPFR knockdown in Griddle-2 in larvae fed on sucrose only (top row) or starved (bottom row). Left panel calcium responses of Griddle-2, for each trial Lower right panel: mean calcium response of Griddle-2 over time. The green dashed line corresponds to stimulus onset. Lower top panel: mean calcium response averaged during the stimulus. White line represents the mean, white dot represents the median. sNPFR knockdown does not influence the stimulus-induced activity of Griddle-2 in larvae fed on sucrose only or starved (n = 10 larvae, 3 trials per larva).

